# Identification of lipid specificity in membrane organizing protein complexes

**DOI:** 10.64898/2026.07.24.740489

**Authors:** Katelyn C. Cook, Karen Palacio-Rodriguez, Kristin Böhlig, H. Mathilda Lennartz, Swantje Lenz, Andrej Shevchenko, Alf Honigmann, Gerhard Hummer, Alexander von Appen, André Nadler

## Abstract

Organelle membrane identity is encoded by protein and lipid composition^1^. Human cells produce thousands of distinct lipid species for this purpose^2,3^. Yet, the precise molecular functions of most lipids remain unknown, owing to limited methods for *in situ* investigations of individual lipid species. Here, we use minimally modified bifunctional lipid probes to detect and functionally characterize lipid-protein interactions across time and subcellular compartments. We leverage this approach to map time-resolved protein interactomes of individual lipids representing major membrane lipid classes as they are transported through the organelle system of human cells. We identify hundreds of unique lipid-protein interaction candidates, measure lipid-induced protein abundance changes, and distinguish between directly and indirectly interacting subunits of associated protein complexes. Informed by this multimodal dataset, we interrogate the lipid specificity of membrane organizing complexes, in particular the selective co-association of phosphatidylethanolamine with the ancestral subunits of the nuclear pore (NDC1) and MICOS (MIC60) complexes. Using super-resolution microscopy and molecular dynamics simulations, we demonstrate that phosphatidylethanolamine stabilizes both NDC1 and MIC60 in highly curved membrane nanodomains, indicating that major membrane organizing complexes are positioned by selective lipid-protein interactions. Altogether, our experimental strategy provides a blueprint for discovering and characterizing cellular functions of individual lipids.

---

Membranes compartmentalize and coordinate cellular biochemistry. Their structural building blocks are lipids and proteins, often present in a nearly equal mass ratio. Human cells, in particular, produce ∼2000 molecularly distinct lipid species, which often vary by only a few atoms, and encode ∼5,000 integral membrane proteins in their genome^1,3–6^. Our understanding of membrane protein function greatly exceeds that of lipid function, yet lipids outnumber proteins by up to 100-fold in organelle membranes and their dysregulation causes numerous monogenic and complex diseases. This is due to two main factors: (i) Unlike proteins, which often act as molecular machines that can be studied in isolation, lipids always exert their functions as components of molecular assemblies^3,7^; (ii) the classical experimental blueprint of genetic perturbations, structural analysis, and fluorescent fusions is largely unsuitable for lipid cell biology. Mechanistic analysis of lipid function in cells is complicated as lipid structure is not directly encoded in the genome, lipids are rapidly moved between organelle membranes and metabolically inter-converted^8–13^, and finally, even minimal structural differences can significantly impact lipid behavior^14–17^. Addressing these challenges inherent to lipid biology is necessary to answer key open questions in particular with regard to the organizing principles of organelle membrane structure and function.

Several notable advances in chemical biology have begun to address this methodological barrier^18,19^. Combining modified enzymes and biorthogonal chemistry has enabled live-cell measurements of local lipid metabolism and transport^20–24^, lipid biosensors track the dynamics of endogenous lipids^25–27^, and caged lipid probes allow for quantitative lipid biochemistry in living cells^28–30^. Microscopy^31^ and mass spectrometry (MS; reviewed in^32^) developments facilitate increasingly accurate observations of lipid spatial and chemical organization^33–36^ as well as biochemically stable lipid-protein complexes^37,38^. Bifunctional (cross-linkable and clickable) lipid probes were developed to capture a greater spectrum of lipid-protein interactions with minimal disruption of lipid structure^39–46^. We recently introduced an approach using bifunctional lipids to quantify species-specific lipid localization, metabolism, and transport between organelle membranes^14,15,47,48^. However, a comprehensive workflow for discovering the biological functions of individual lipid species is still needed, as it must simultaneously: (i) address small molecular differences between species; (ii) resolve rapid lipid redistribution between membranes; (iii) account for lipid metabolism and (iv) lipid-induced protein abundance changes; and (v) capture both direct lipid-protein interactions and multi-component lipid-protein assemblies in the native membrane environment (**Figure 1A**).

**Figure 1:**
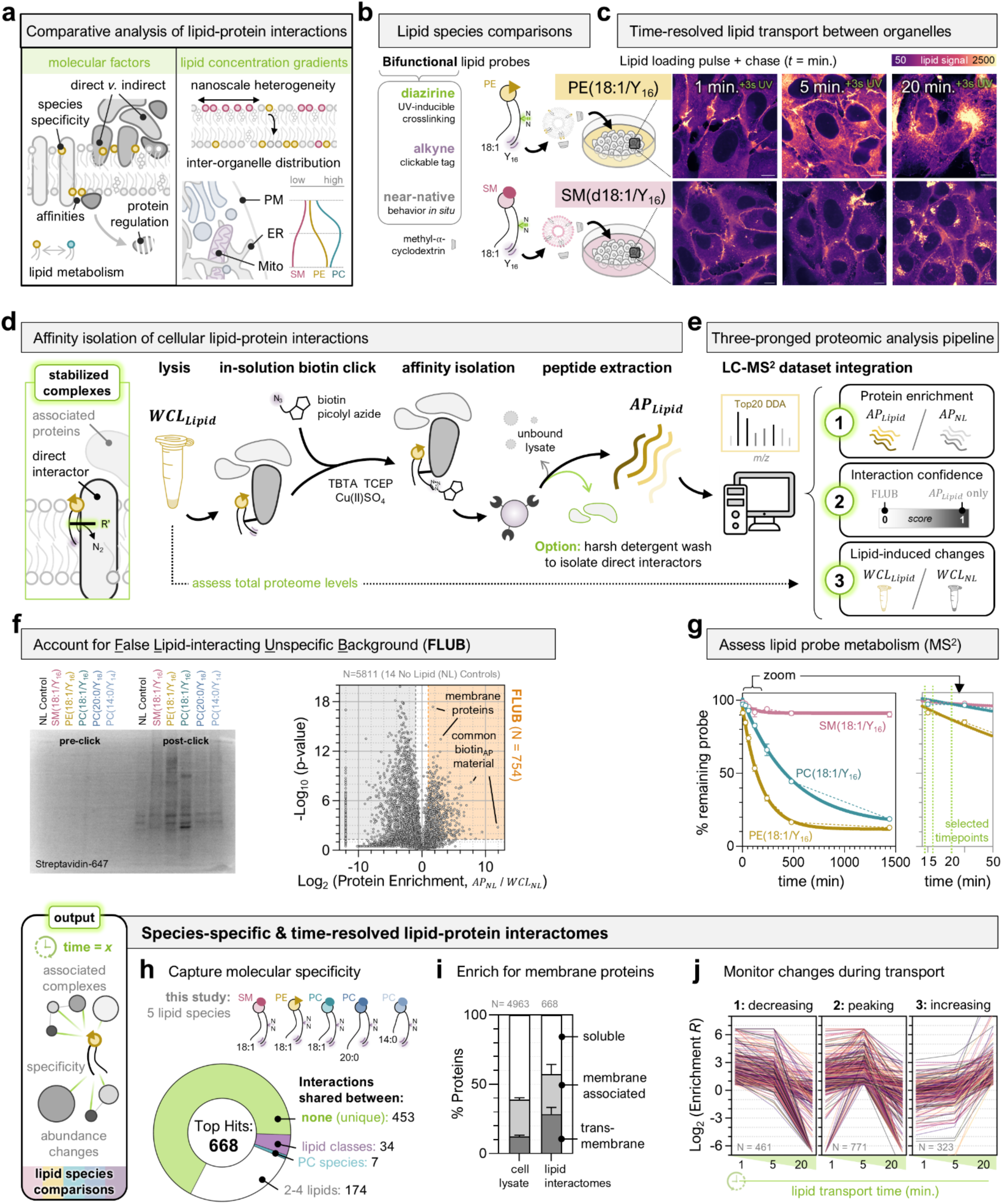
Bifunctional phospholipid probes enable lipid-specific protein interactomics *in situ*. **A)** Detection and interpretation of lipid-protein interactions in cellular membranes are influenced by: 1) lipid species specificity, often confounded by rapid metabolic inter-conversions, 2) direct interactions versus indirect or associated protein complexes, 3) differing affinities or interaction types (e.g., binding pocket versus membrane environment), 4) lipid bioactivity resulting in changing abundances of protein interactors, and 5) different and rapidly changing subcellular lipid densities between organelles, membrane subdomains, and membrane leaflets. **B)** Bifunctional phospholipid probes contain a mid-chain diazirine (UV activatable crosslinker) and terminal alkyne (click chemistry tag) on one fatty acid tail. Species diversity of native lipids is captured by chemical synthesis of probes that vary in headgroup, backbone, and acyl chain structure to mirror native lipids. Probe notation is *Headgroup(acyl chain sn-1/acyl chain sn-2)*, with Y_x_ denoting the bifunctional chain and *x* = number of carbons. Two examples are shown as cartoons: SM(d18:1/Y_16_ (*pink*) and PE(18:1/Y_16_) (*yellow*). Target probes are incorporated into liposomes prior to cellular loading. **C)** Live cell cultures receive lipid probes via a methyl-*a*-cyclodextrin mediated exchange pulse that loads lipids into the outer leaflet of the plasma membrane (pulse = 45sec. for the 1min. timepoint; 3.5min. for the 5 and 20min. timepoints). Probes commence immediate intracellular trafficking throughout the organelle system and are locked in place by rapid UV irradiation (3sec.) at the desired timepoint (e.g., 1, 5, 20 minutes post-pulse). Lipid localization is visualized by fluorescent lipid imaging^14^ (U2OS cells; scale bars = 10µm). **D)** Biochemical workflow for the lipid-protein affinity pulldown (LP-AP). The protocol is optimized to retain both covalently bound and associated membrane proteins following UV-induced diazirine crosslinking (*left*); associated proteins can later be separated by a harsh wash (high % detergent) prior to elution (*right*). A portion of the input lysate (10%; *WCL_Lipid_*) is set aside for whole-cell proteome analysis to quantify protein abundances in the starting sample; the remaining sample (*AP_Lipid_*) undergoes copper-mediated biotin click, streptavidin bead isolation, and hydrophobic peptide preparation for MS analysis. Paired “no lipid controls” (NL) are included with every sample acquisition to account for the variability inherent to membrane biochemistry. **E)** Three proteomic datasets are integrated to identify top-candidate hits from pulldown (AP) and whole-cell lysate (WCL) samples. Ultimately, every detected protein receives an enrichment ratio (*1*) that is scaled by total protein abundance (*3*), comparing the lipid to NL control samples. Confidence scores are generated from analysis of the NL control AP and WCL datasets (*2;* see *FLUB* in **F**). **F)** *Left,* Western blot of whole-cell lysates pre- and post-biotin click, stained for streptavidin (NL control versus five lipid pulldowns; sample type labeled at *top*). Streptavidin signal in the NL control indicates material non-specifically binding to biotin following the click reaction, shared across all samples. *Right,* Volcano plot of MS quantification of proteins enriched by LP-AP in the NL controls (N= 5811; 7 biological x 2 technical replicates), plotted with protein enrichment (*x-*axis; Log_2_(AP_NL_/WCL_NL_)) against significance (*y-*axis; p-value). Proteins significantly enriched by the protocol are termed FLUB. All proteins in the left grey area (AP_NL_/WCL_NL_ ≤1.5, p ≤0.05) are properly depleted by the protocol. See also **Extended Data Figure 3** for full FLUB analysis. **G)** Ultra high-resolution lipid mass spectrometry quantification of bifunctional probe abundance (*y*-axis) over time (*x*-axis). Each point is the mean, error bars are standard deviation, and solid lines represent the mono-exponential decay (least squares fit). Data reproduced from Iglesias-Artola *et al.*^14^ **H-J)** Summary of the top-candidate lipid-protein interactions identified in this study (N=668 proteins; R≥1.5, p≤0.05, S≥0.5; N=3 biological x 2 technical replicates for 5 lipid species), shown as **H)** a donut plot of the number of top hits shared between lipids (key at *right*), **I)** a stacked bar plot of proteins annotated to be transmembrane, peripheral/membrane associated, or soluble, comparing input cell lysates to lipid interactomes (bars are mean; error bars are standard deviation across lipid conditions), and **J)** line graphs of lipid-protein interactions across time, clustered by decreasing, peaking, or increasing trends (each line is a protein, MS quantification of mean protein abundance at 1, 5, 20 minutes post-lipid loading for PE pulldowns is exemplarily shown).

Here, we integrate lipid imaging with lipid-protein affinity capture to systematically identify the interactions of lipid species with individual proteins and membrane-associated protein complexes during transport through the organelle network, creating an experimental pipeline for mechanistic lipid cell biology. We map the time-resolved protein interactomes of individual phospholipid species as they are trafficked through the organelle membrane network of human cells. We find that each lipid probe has a unique protein interactome, with an average of 68% of protein interactors specific for a single lipid. By accounting for lipid-induced changes to protein levels and variable interaction specificities, we distinguish between lipid biological activity and *bona fide* lipid-protein interactions. To explore the close connections between lipid-protein interactions and membrane organization and shape, we further investigate the selective interactions between the mitochondrial intermembrane space bridging (MIB; contains MICOS) and nuclear pore (NPC) complexes with phosphatidylethanolamine (PE). Using STED super-resolution microscopy and molecular dynamics (MD) simulations, we show that the core transmembrane components of both major membrane organizing complexes, MIC60 and NDC1, are stabilized in highly curved membrane segments by interactions with PE at the inner bilayer leaflet.

## Time-resolved lipid-protein interactomes

We developed an experimental strategy for comparative studies of species-level lipid cell biology. Our approach relies on a pulse-chase strategy using minimally-modified photoactivatable (diazirine) and clickable (alkyne) bifunctional phospholipids (**Figure 1B**), which recapitulate the behavior of native lipids^14^. Lipids are delivered to the outer leaflet of the plasma membrane in a brief loading pulse (∼1 minute) using an alpha-methyl-cyclodextrin mediated exchange reaction with donor vesicles^14,49^. Lipid transport through the organelle system commences immediately. After the desired chase time (e.g., 0-16 minutes), a rapid UV irradiation (3 seconds) covalently crosslinks the lipid probes to adjacent proteins via diazirine photoactivation^43^ (**Figure 1C; Extended Data Figure 1**). For lipid imaging, we immediately chemically fix, permeabilize, and attach a fluorophore to the alkyne group in the fatty acid moiety by click chemistry^14^. For proteomics profiling, we freeze and then lyse cell pellets, attach a biotin tag using in-solution click chemistry, and perform streptavidin-mediated affinity isolation and peptide extraction for MS analysis (**Figure 1D**; **Extended Data Figure 2**). We optimized the lipid-protein affinity pulldown (LP-AP) workflow to retain both the covalently crosslinked interactors as well as associated protein complex components to retain biological context. Direct and indirect interactors are distinguished experimentally by treatment with a strong detergent prior to elution. Proteomics data was acquired using optimized data-dependent LC-MS/MS (see **Methods**).

Our LP-AP data analysis pipeline is designed to systematically account for both biological and technical variability in detecting, quantifying, and interpreting crosslinked lipid-protein complexes. To this end, we integrate four MS datasets to generate each lipid interactome (**Figure 1E**). To distinguish high-confidence lipid-protein interactions from non-specific pulldown background, we first compare the actual lipid pulldown sample containing interacting proteins (*AP_Lipid_*) with its paired “no lipid” (NL) control pulldown sample (*AP_NL_*), which is not lipid-treated but otherwise handled the same (**Figure 1F**). We use these datasets to calculate a pulldown enrichment ratio 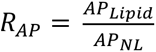 for each detected protein. Next, we account for lipid-induced changes to detected protein abundances by acquiring total proteome quantifications from the whole-cell lysates prior to pulldown (*WCL_Lipid_* and *WCL_NL_*), which also contain crosslinked lipid-protein complexes. From these datasets, we calculate a relative abundance ratio 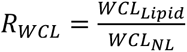 for each protein that reflects changes to detectable peptide MS intensities following lipid treatment and UV-photoactivation (**Supplemental Table 6**). Since both true lipid-protein interactions and underlying changes to detectable peptide abundances would affect interaction quantification in our assay, we integrate both ratios into a scaled enrichment value 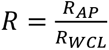 for each protein. The scaled enrichment value *R* is particularly useful in two cases: i) for low abundance proteins that do not meet MS detection criteria in the whole-cell proteome (a common problem for membrane-associated proteins^50–52^) yet are well-detected in the lipid pulldown sample; ii) for proteins whose MS detection is confounded by lipid crosslinks, as the crosslinked lipid-peptide species exhibit different biochemical and LC-MS/MS behavior (discussed in **Supplemental Information**).

We further score interaction confidence by pooling the information from all ten “no lipid” control datasets (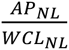; 5811 proteins). We define proteins that are non-specifically and significantly enriched by our protocol as the false lipid-interacting <u>u</u>nspecific <u>b</u>ackground (FLUB; approximately 13% of quantified proteins) (**Figure 1F; Supplemental Table 1**). Conceptually, this is similar to the CRAPome for protein-protein interactome scoring^53–55^. Indeed, we find that ∼30% of FLUB proteins are frequently reported in the CRAPome database (>10% of biotin affinity pulldowns), while the remaining proteins largely map to insoluble membrane compartments (**Extended Data Figure 3**). Control datasets are also used to assign a confidence score (*S*) to candidate interactors, ranging from 0 to 1. FLUB is 0, proteins that were depleted by the pulldown are 0<*S*<1, and the highest confidence interactors that are never detected in a control pulldown receive *S*=1. We ultimately define “top hits” likely to reflect *bona fide* lipid-protein interactions as proteins with an enrichment ratio *R* ≥1.5, p-value ≤0.05 (Benjamini-Hochberg corrected), and confidence score *S* ≥0.5.

Crosslinking probability is a function of lipid-protein affinities as well as lipid levels in an organelle membrane. We address the latter by combining the MS analysis with lipid imaging to track changing subcellular localization (**Figure 1C**) and lipid MS measurements to track probe metabolic inter-conversions (**Figure 1G**). Using this information, we can directly compare individual phospholipid species in parallel experiments—rather than pan-lipid labeling by metabolic precursors—by choosing the most informative timepoints for proteomic analysis: when the majority (∼90%) of the bifunctional probes remain our target species and the unique organelle distributions of different lipids allow for effective interactome interpretation. Altogether, our time-resolved imaging-proteomics approach identifies lipid-protein interactomes that are species-specific (**Figure 1H**), capture the target membrane proteome (**Figure 1I**), and reflect lipid movement across chemical and physical space (**Figure 1J**).

## Species-selective lipid-protein networks

We chose five probes representing the main structural glycerophospholipid classes in U2OS cells: phosphatidylcholine (PC; ∼33% of all lipids) and phosphatidylethanolamine (PE; ∼20%), as well as sphingomyelin as the primary sphingolipid class (∼6%)^14^. We compared common (PC(18:1/Y_16_), PE(18:1/Y_16_), SM(d18:1/Y_16_)) and rare lipid species (PC(20:0/Y_16_), PC(14:0/Y_14_); ∼0.0001%) designed to isolate the effects of headgroup variation versus chain length and saturation. We first performed a comparative analysis of all probes at 20 minutes post-lipid delivery, a timepoint when limited probe metabolism has occurred (<2-13%) (**Figure 1H**). Lipid imaging confirmed that probes were incorporated to a similar degree and exhibit unique organelle localization profiles (**Extended Data Figure 1, Figure 2A**). Specifically, SM has a pronounced localization at the plasma membrane and vesicular structures, whereas PE is primarily localized to ER and mitochondria, and the PC species showed localization profiles between these two extremes. To account for the temporal trends of lipid-protein interactions during lipid transport, we also compared earlier timepoints (1 and 5 minutes) for SM and PE probes, which have very different steady-state distributions and trafficking mechanisms (**Figure 3**).

**Figure 2.**
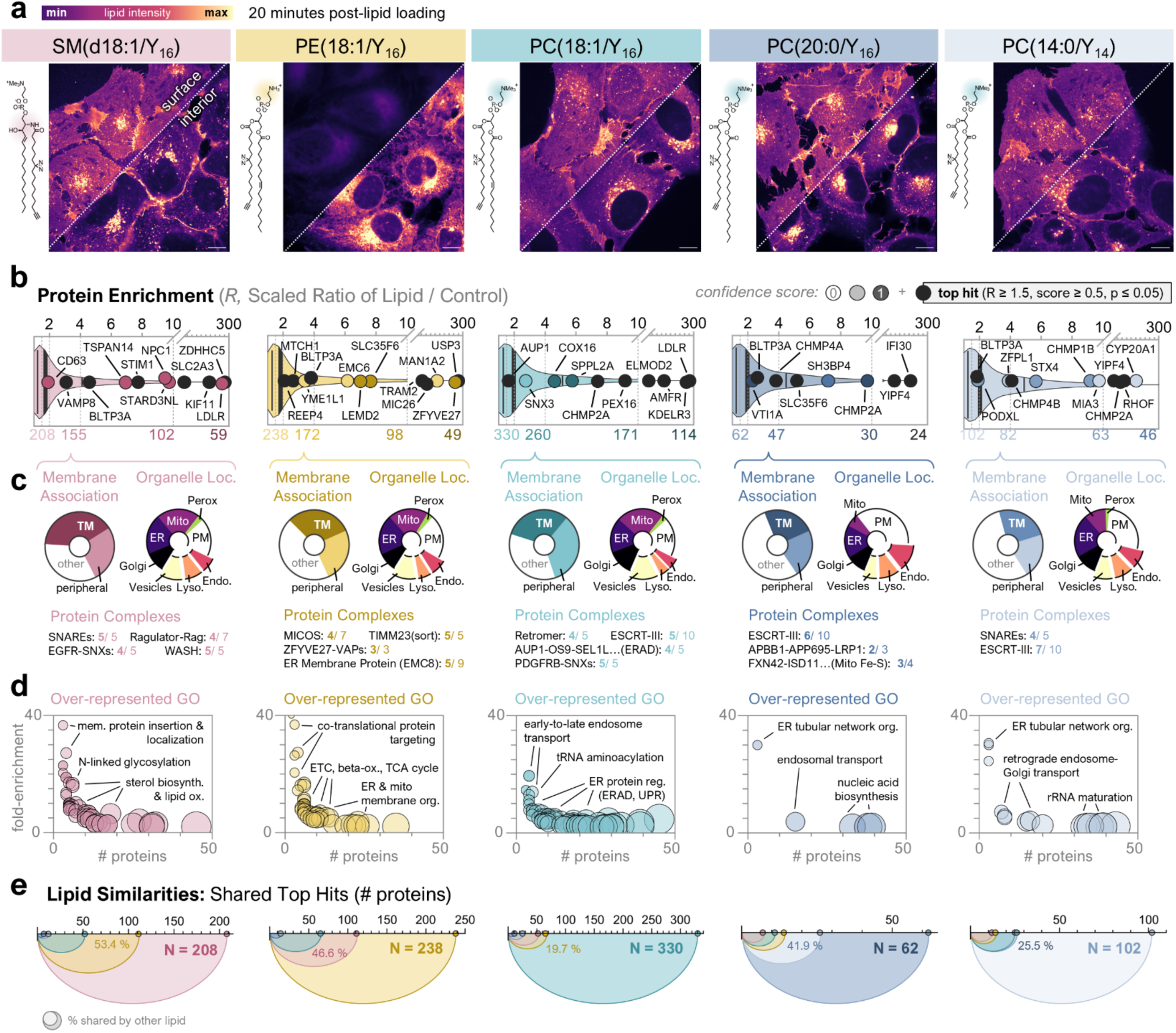
Membrane lipid species engage unique protein networks. **A)** Representative images of U2OS cells at 20 minutes post-lipid pulse, showing a basal surface (plasma membrane) and interior *z*-slice for each field of view (separated diagonally by dotted white line; scale bars = 10µm). Chemical structure is at *left* and probe notation at *top*, sorted from *left* to *right* by major membrane lipid classes (SM(d18:1/Y_16_ (*pink*), PE(18:1/Y_16_) (*yellow*), PC(18:1/Y_16_) (*teal*)) and rare PC species (PC(20:0/Y_16_) (*dark blue*), PC 14:0/Y_16_ (*light blue*)). All images are shown on the same intensity scale (*top left*) for comparison. **B)** Protein interactomes for each lipid (corresponding to images in **A**) summarized in violin plots of the scaled enrichment ratio (*R; x*-axis). Representative proteins are shown as labeled points, colored by increasing interaction confidence (*S* and p-value; key at *upper right*). The number of top hits (*R*≥1.5, p≤0.05, S≥0.5) passing increasing enrichment thresholds (R≥*x* at dotted lines) is indicated *below*. **C)** Feature annotations for top hits (*R*≥1.5). Pie charts show membrane association (*left*; TM= transmembrane) or organelle assignment (*right*). CORUM^56^ protein complexes with >50% of members represented (N = enriched / total). **D)** Gene ontology terms from a PAN-GO^57^ overrepresentation analysis (Fisher’s exact test; enrichment >2, FDR< 1%, p≤ 0.05) are shown as a multi-variable plot (fold-enrichment to human proteome on *y*-axis, # top hit proteins for each term on *x*-axis; node size scales with # proteins in each term). Three of the most prevalent parent terms are labeled on each graph. **E)** Venn Diagram comparison of top hits for each lipid displaying the fraction of shared hits with the other four lipids. The total number of hits is on the *x*-axis and colors are indicative of individual lipids as defined in **A**.

**Figure 3.**
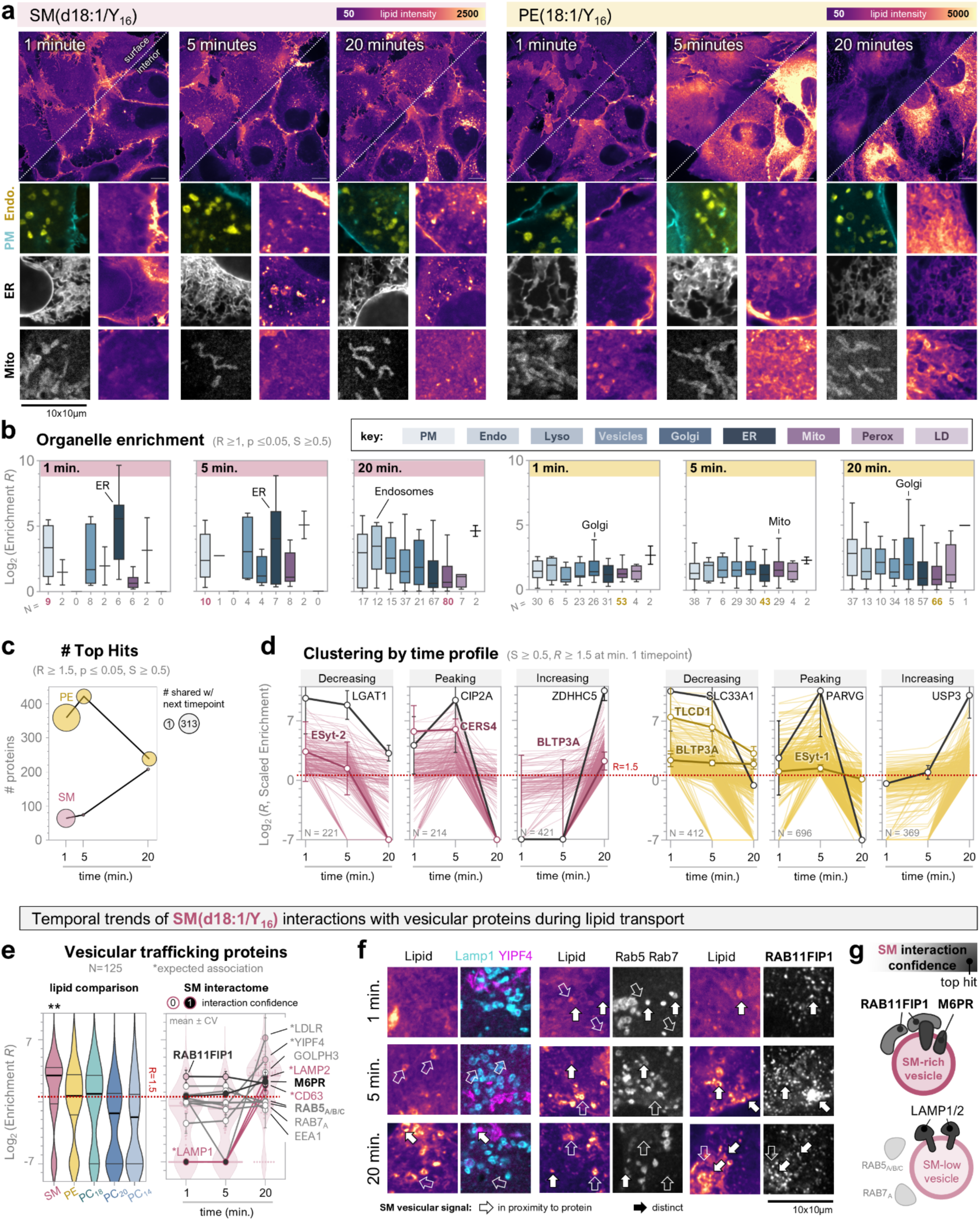
Lipid-protein interactions change over time during inter-organelle lipid transport. **A)** Representative images of U2OS cells at 1, 5, and 20 minutes after loading SM(d18:1/Y_16_) (*left*; pink) or PE(18:1/Y_16_) (*right*; yellow). At top is a larger field of view showing a basal surface (plasma membrane) and interior *z*-slice (separated diagonally by dotted white line; scale bars = 10µm). Below are three 10×10µm regions from each timepoint showing organelle co-stains (*left*) and the corresponding lipid channel (*right*). Antibodies against endogenous proteins were used for endosome and mitochondria co-stains (see **Methods**), ER is labeled by HaloTag-Sec61β, and plasma membrane (PM) by SNAP-PDGFR_TM_ (see also **Extended Data Figure 1**). **B)** Box-and-whisker plots showing organelle-specific protein enrichment by LP-AP (*y*-axis; Log_2_) at each timepoint corresponding to the above images in **A** (1, 5, 20 minutes left-to-right; *pink* = SM, *yellow* = PE). The organelle color *key* is at top, with endomembrane organelles in blue and others in purple. The top organelle for each timepoint is labeled. Whiskers are Tukey distribution with line at median; N values *below* indicate the number of proteins for each organelle at that timepoint. Proteins plotted here are interactor candidates with a relaxed filter of *R* ≥1 (still S ≥0.5, p ≤0.05; N = 3 biological x 2 technical replicates) to include all enriched proteins. **C)** The total number of top hits for each lipid plotted across time (*x-*axis). Node size corresponds to the number of protein interactions that are shared with the following timepoint (within the same lipid). **D)** Interactor candidates clustered by decreasing, peaking at 5 minutes, or decreasing enrichment (*y*) across time (*x*). Each line is a protein (N value indicated at bottom left of each plot). The top representative protein for each cluster is labeled and colored black with node at mean and error bars ±%CV. **E)** Plot of vesicular trafficking protein enrichment by different lipid species at 20 minutes (*left*) and SM across time (*right*). Lipid comparisons are shown as violin plots of N=125 proteins annotated to be involved in vesicular trafficking (bold line at median, thin lines at quartiles; **p<0.01 by two-way ANOVA). For the SM interactomes across time, each line is a protein with node at mean and error bars ± %CV (pink indicates proteins with previously reported sphingolipid associations*). The violin plots in the background show the total distribution of N=125 vesicular trafficking proteins (see also **Supplemental Tables 3-4**) **F)** 10×10µm regions of U2OS cells showing vesicular SM localization at 1, 5, and 20 minutes post-probe addition. The lipid signal is at *left* and the corresponding protein co-stains at *right* (antibodies against endogenous proteins). All proteins are represented in the graph at **E.** White arrows indicate vesicular SM that does not co-localize with the protein co-stain; green indicates co-localization (often proximal subdomains). **G)** Cartoon summarizing the protein components involved in inter-organelle SM transport, as suggested by this study (*red*) and known previously (*black*).

We identified hundreds of putative lipid-protein interactions at 20 minutes post-probe addition, including 668 top-candidate interactors (*R* ≥1.5, p ≤0.05, *S* ≥0.5). Among these, 68% were uniquely detected for a single lipid probe (**Figure 1H**, **Figure 2E, Extended Data Figure 4**). The more abundant membrane phospholipids (SM, PE, PC(18:1/Y_16_)) have a higher number of top hits compared to rare PC species, and a higher fraction of transmembrane proteins (**Figure 2B-C**). Very few hits are shared between lipid classes (34) and PC species (6), respectively. The limited overlap between lipid-protein interactomes strongly suggests that interaction candidates represent molecularly distinct lipid-protein assemblies rather than the general organelle membrane proteome. This is further supported by the observation that lipids with similar subcellular distribution (e.g., PC species; **Figure 2A; Extended Data Figure 1**) still enrich for distinct protein networks (**Figure 2C-D**). We note that there is an abundance of biology to explore here, expanding upon our current knowledge of lipid cell biology and species-selective lipid function^7,8,10,11,26,35,58–61^. We expect our datasets to serve as a comprehensive resource for future studies in lipid and membrane biology (**Supplemental Tables 2-3**). Here, we limit the discussion of detected lipid-protein interactions to a few selected examples that illustrate potential interactome interpretations and effective selection of candidates for functional validation.

For SM(d18:1/Y_16_), we find enrichment of proteins involved in vesicular trafficking (e.g., CD63, KIF11, VAMP3/8), cholesterol transport (STARD3NL, the sphingosine interactor NPC1^42,62^), glycosylation (ALG2, C1GALT1C1, B3GAT3, CAMLG, several OST complex components), protein lipidation (the top hit ZDHHHC5), as well as GPI-anchored proteins (MELTF, TSPAN14) (**Figure 2B-D, Supplemental Table 3**). For PE(18:1/Y_16_), there are many proteins involved in ER and inner mitochondrial membrane organization, including components of the MICOS, TIMM, and nuclear pore complexes, ER network proteins (REEPs, ZFYVE27, VAPA, EMC components), MTCH1 which was recently suggested to have scramblase activity^63^, and the mitochondrial transmembrane protease YME1L1 (**Figure 2B**), which functions in mitochondrial homeostasis and is directly regulated by cellular PE levels^64,65^. For all PC probes, we enrich for general membrane remodeling proteins including the ESCRT-III machinery, where CHMP2A is a top hit for all three species. PC(18:1/Y_16_), an analog of one of the most common lipid species in human cells, has the most interactor candidates, among them components of protein quality control (e.g., AUP1, AMFR) and tRNA aminoacylation machinery (**Figure 2C-D**).

Among the shared top hits (2-4 lipids), we observe many proteins that have been previously shown to either directly regulate, or be regulated by, membrane lipids. First, the bridge-like transfer protein BLTP3A is a top hit for every lipid except PC(18:1/Y_16_), where it is only just below the threshold (*R =* 1.49) (**Figure 2B**). SM and PE(18:1/Y_16_) exhibit the highest enrichment for BLTP3A, and interestingly these lipids also enrich for the ER-plasma membrane contact site acyltransferase TLCD1, a regulator of bridge-like transfer protein BLTP2 that was previously shown to influence subcellular PE levels^59,66–70^. Only seven other proteins are top hits for four-out-of-five lipids, including the plasma membrane glucose transporter SLC2A3 (GLUT3), and the secreted lysosomal glycoprotein precursor prosaposin (PSAP), which binds sphingolipids and is a genetic determinant of sphingolipid storage disorders^71^. The three structural phospholipids share several other compelling transmembrane interactor candidates, including the ER translocator TRAM2 (functionally influenced by sphingolipid binding^72^), and the palmitoylated GTPase activating protein (GAP) ELMOD2 that regulates lipid droplet, mitochondria, and ciliary membrane biogenesis^73,74^ (**Extended Data Figure 4F**).

## SM enriches for non-canonical vesicles

We next mapped changes in lipid-protein interactions during lipid transport from the plasma membrane to intracellular organelles (**Figure 3, Supplemental Table 3**) for SM(d18:1/Y_16_) and PE(18:1/Y_16_). Specifically, we compared interactomes from a 1-minute timepoint, where both lipids still exhibit strong plasma membrane localization, with an intermediate 5-minute timepoint and the 20-minute timepoint, which reflects the steady-state distribution (**Figure 3A, Extended Data Figure 1**). The number and type of interactions change as lipids are transported (**Figure 3C**). At 1 minute, the primary organelle enrichment for SM is at the plasma membrane, vesicles, and ER, reflecting the imaging results, while PE interaction candidates appear to be distributed throughout the organelle system. At 20 minutes the primary organelle enrichment for SM are non-ER organelles of the secretory pathway, a trend that is not observed for PE (**Figure 3B**). PE has a significant number (313) of shared top hits between timepoints, reflecting near-immediate incorporation throughout the ER and mitochondrial networks (**Figure 3B-C**). In contrast, SM has only 1 shared top hit from 1 to 20 minutes, which likely reflects the accumulation in specific vesicular compartments (**Figure 3F**, **Extended Data Figure 5**).

We observe temporal trends in lipid-protein interactions that may underlie lipid-specific transport mechanisms. For example, PE enriches for distinct membrane contact site proteins and lipid transfer proteins at all timepoints, reflecting that PE is primarily transported by non-vesicular routes^14,61,68,75^. This includes BLTP3A, TLCD1 (regulator of BLTP2), and the ER-plasma membrane tethering transporter E-Syt1 (**Figure 3D, Extended Data Figure 4G-H**). In contrast, SM enriches for BLTP3A only at 20 minutes, for TLCD1 only at 1 and 20 minutes, and for E-Syt2 (1-5 minutes) rather than E-Syt1. In addition, at 1 minute the most highly enriched SM interactor candidates are ER proteins involved in sphingolipid biosynthesis (e.g., ceramide synthases CERS4/5/6 and the sphingolipid desaturase FADS3; **Figure 3B & D**), while by 20 minutes vesicular trafficking proteins are over-represented. This includes proposed sphingolipid interactors like LAMP1/2^9,26^, NPC1^42,62^, LDLR, the tetraspanins CD63 and TSPAN14^76–78^, and many Golgi proteins^62,79^ (e.g., SACM1L, TMEM230, YIPF4, GOLPH3), which are more enriched for SM compared to all other lipids (**Extended Data Figure 5A**). The temporality of these interactions captures the shift from non-vesicular lipid internalization (plasma membrane to ER, <5 minutes) to vesicular anterograde cycling (back to plasma membrane, >15 minutes) that we previously showed for SM^14^.

Interestingly, at all timepoints SM exhibited higher enrichment of non-canonical RABs and RAB effectors (e.g., RAB11FIP1, RABAC1, RABL3, RAB39B) than proteins that are typically associated with endosomes (e.g., RAB5A/B/C, EEA1, RAB7, RAB11A/B, SNX1), lysosomes (e.g., LAMP1/2, CD63), or Golgi (e.g., RAB1B, YIPF4, GOLPH3) (**Figure 3E, Extended Data Figure 5B**). This observation intrigued us, as it suggested that the SM pulldown enriched for a specialized vesicular compartment involved in SM transport. To validate the time- and protein-dependent trends in our interactome data, we performed immunofluorescent imaging experiments of SM(d18:1/Y_16_) with antibodies against vesicular interactor candidates at 1, 5, and 20 minutes post-lipid loading. Overall, lipid-protein co-localization confirmed the detected interactions. SM-rich vesicles are visible at all timepoints, growing brighter over time as lipid accumulates in the compartment (**Figure 3F**), in-line with increased number of vesicular protein interactors by 20 minutes. SM-rich vesicles were infrequently co-labeled by RAB5/7 at all timepoints, while LAMP1 and/or Golgi (labeled by GM130 or YIPF4) co-localization was observed primarily at 20 minutes (**Figure 3F-G, Extended Data Figure 5C-D**). However, the brightest SM vesicles were nearly always lacking these canonical endocytic markers. We thus selected a protein from the highly enriched non-canonical candidates: RAB11FIP1, the only one enriched by SM at all timepoints (**Figure 3E**). In this case, nearly all SM-enriched vesicles exhibited close proximity to RAB11FIP1 puncta (**Figure 3F-G**).

Given the rapid and sustained enrichment of the RAB11FIP1 by SM, it is possible that RAB11FIP1 is an upstream assembly factor for SM sorting machinery in vesicular compartments. RAB11FIP1 has indeed been reported to regulate endosomal recycling, acting as a coupling factor for RABs like RAB11^80^, and SM transport requires entry into anterograde transport vesicles^9,14,26^. However, it is unlikely that RAB11FIP1 directly senses SM levels within the membrane, as it is a cytosolic protein while SM is expected to asymmetrically localize to the luminal leaflet of the vesicular bilayer^20,35^. To test this, we performed a SM pulldown experiment using a harsh detergent wash (0.2-1% SDS) of the beads to remove indirect interactors before elution (e.g., non-crosslinked peripheral subunits of the lipid-protein complex), selecting for direct protein interactors (e.g., covalently crosslinked lipid-protein conjugates that are retained following 1% SDS; **Supplemental Table 5**). RAB11FIP1, RAB5/7, and most other cytosolic RABs did not survive SDS treatment, indicating that they are associated rather than direct interactors. In contrast, other interactions with vesicular proteins were maintained, including LAMP2, the transmembrane RAB regulator RABAC1 (also known as YIP3), the SNAREs VAMP3/8, the ER-endosome contact site sterol transporter STARD3NL, and M6PR, a known regulator of sphingomyelinase and the trans-Golgi cargo-sorting receptor for lysosomal targeting^81,82^ (**Extended Data Figure 5E-F**). RAB11FIP1 could conceivably act in complex with these proteins to build specialized SM sorting compartments. Interestingly, this complex is likely independent of RAB11, as RAB11A and B were well-quantified in our datasets yet not enriched by SM at any timepoint (**Extended Data Figure 5B**). Altogether, these experiments illustrate the utility of our approach for characterizing a transient, protein- and lipid-specific compartment that would be otherwise inaccessible by other methods, leading us to a simple model where RAB11FIP1 coordinates entry of SM into non-canonical vesicles during recycling.

## PE localizes at MICOS and NPC domains

One of the most prevalent trends in our datasets is the over-representation of membrane organizing complexes in the PE(18:1/Y_16_) interactomes (**Figure 2F**). These included the mitochondrial intermembrane bridging complex (MIB; N= 21/23 proteins detected), which contains the mitochondrial contact site and cristae organizing system (MICOS), and the inner ring of the nuclear pore complex (NPC; N= 29/32 proteins detected) (**Figure 4A-B**), which are conserved across all eukaryotes. Both complexes were more enriched by PE compared to all other lipids tested, especially for the core ancestral transmembrane subunits of each complex, MIC60^83,84^ and NDC1^85–87^, which were top hits only for PE (**Figure 4C-D, Extended Data Figure 6**). PE lipids have conical shape, an intrinsic property that enables bilayer deformations required for membrane curvature *in vitro*^88–91^. The MICOS and nuclear pore complexes are essential residents of some of the most curved membrane domains in eukaryotic cells, with mean radius of curvature *r*=10-30 and 30-50 nm for mitochondrial cristae junctions and the nuclear pore neck, respectively^92–97^. As in-cell evidence for PE species influencing protein localization to curved membrane domains has remained elusive, the PE-MICOS and -NPC interactions were particularly compelling candidates for studying the organizing principles of lipid-protein assemblies responsible for shaping organelle membranes.

**Figure 4.**
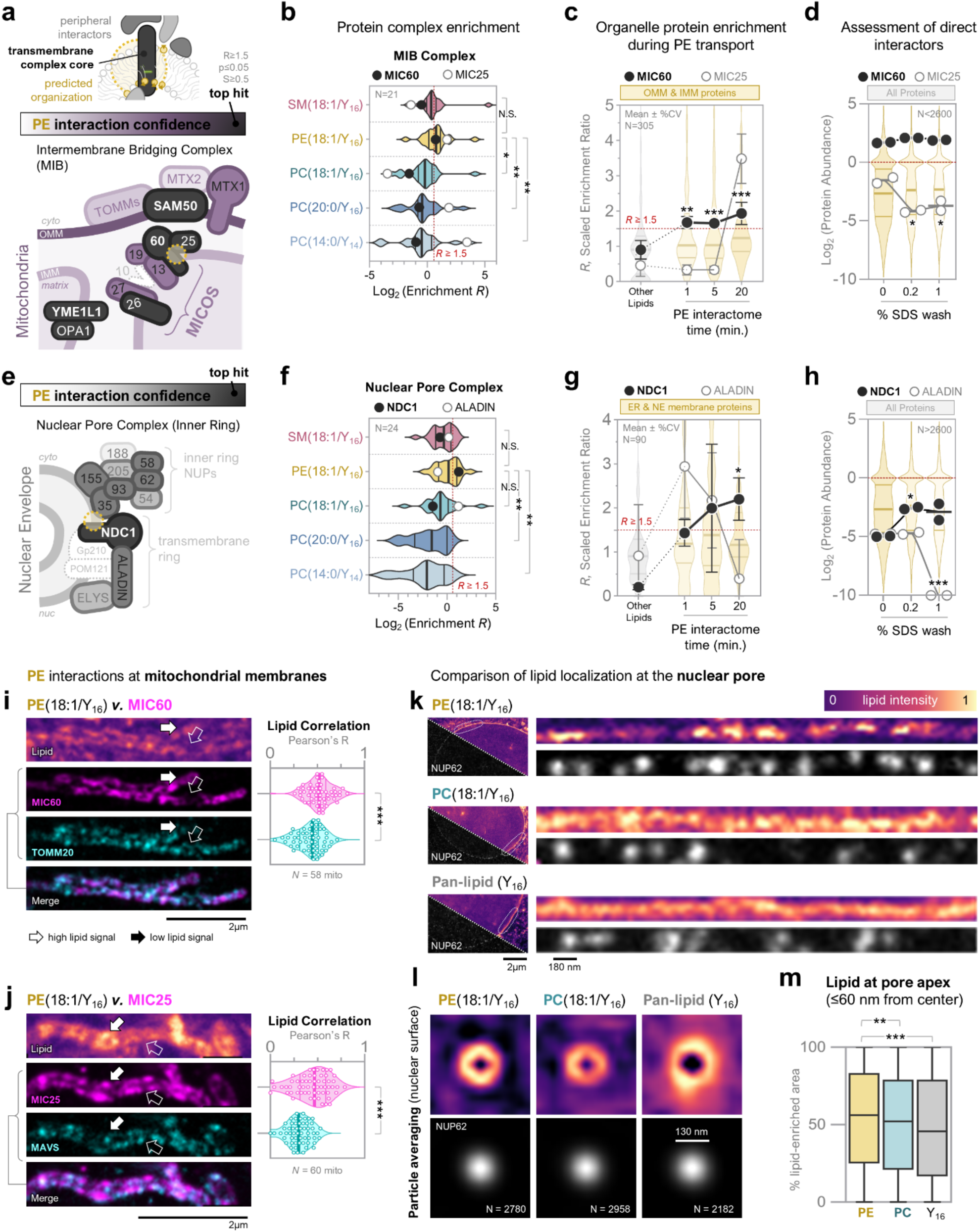
Phosphatidylethanolamine (PE) interacts with MIC60 at mitochondrial cristae and NDC1 at nuclear pores. **A)** Components of the mitochondrial intermembrane bridging (MIB) complex detected in this study, colored by increasing interaction confidence with PE(18:1/Y_16_) from interactome data (Figure 2**-3**; protein color key *above*, with cartoon representation of predicted interaction topologies for candidate PE interactions, whereby darkest color indicates top hits (R≥1.5, S≥0.5, p≤0.05). MIC60, MIC25/26, YME1L1, OPA1, and SAM50 are top hits; MIC10 is not detected. **B)** Comparison of MIB (N=21 proteins) complex enrichment in the lipid species interactomes from this study (20 minutes). The core transmembrane protein MIC60 is shown as black dots, and MIC25 as an example peripheral membrane protein from the same complex is shown as white dots. MIC60 exhibits selective enrichment by PE. Plotted is the scaled enrichment ratio *R* as described in Figure 2, on a Log_2_ scale. Statistical significance was assessed by a nonparametric Kruskal-Wallis multiple comparisons test (*p<0.05, **p<0.01, ***p<0.001; N=3 biological x 2 technical replicates). **C)** MIC60 is an enriched top hit by PE at all timepoints tested, while different trends are observed for the peripheral complex member MIC25 and nearby organellar proteins (outer (OMM) and inner (IMM) mitochondrial membranes; N=305). Plots show the scaled protein enrichment ratio *R* (*y*-axis) at 1, 5, and 20 minutes of PE(18:1/Y_16_) transport compared to the mean enrichment in other lipids (*x*-axis). Individual proteins are displayed as dots with error bars ±CV; violin plots show all other OMM/IMM proteins. Statistical significance was assessed by a Benjamini-Hochberg corrected t-test (*p<0.05, **p<0.01, ***p<0.001; N=3 biological x 2 technical replicates) **D)** The PE-MIC60 interaction is maintained following a harsh detergent wash using SDS while MIC25 is lost. Relative protein abundances quantified by LC-MS/MS are shown (*y*-axis; Log_2_ scale, abundances scaled to mean) following pulldowns with normal (0% SDS) or increasing % SDS washes prior to elution (*x*-axis). Individual replicates for selected proteins are shown as points with line at median, atop violin plots of all other quantified proteins per sample (N>2600). Statistical significance was assessed by Dunnet’s multiple comparisons test to the normal (0% SDS) condition (*p<0.05, **p<0.01, ***p<0.001; N=2 technical replicates). **E)** PE(18:1/Y_16_) interactors at the nuclear pore complex, colored by confidence as in **A.** NDC1 is the only top hit; Gp210 and POM121 are not detected. **F)** Comparison of nuclear pore complex enrichment by the membrane phospholipids in our study, plotted as in **B.** NDC1 (black) is compared to the peripheral protein ALADIN (white); neither were detected in the rare PC species interactomes. **G)** NDC1 is enriched by PE at all timepoints tested, while different trends are observed for ALADIN and other ER and nuclear envelope (NE) membrane proteins (N=90). Plotted as in **C.** **H)** The PE-NDC1 interaction (black dots) is resistant to harsh detergent wash, while the PE-ALADIN interaction (white dots) is lost. Plotted as in **D.** **I-J)** PE lipid imaging in U2OS cells co-labeled with antibodies against endogenous **I) MIC60 and TOMM20** or **J) MIC25 and MAVS**. All four proteins are candidate PE interactors (**Extended Data Figure 6**), with MIC60/MIC25 at the MICOS and TOMM20/MAVS at the outer mitochondrial membrane (OMM). Shown are examples of individual mitochondria at 20 minutes post-PE delivery (150X, single z-slice, SoRA spinning disc; scale bars below). Graphs at *right* display quantification of Pearson’s correlation of lipid versus each protein along the mitochondrial length (N≥58 mitochondria). PE enriches at MICOS membrane domains. Statistical significance by paired Wilcoxon test (*p<0.05, **p<0.01, ***p<0.001). **K)** PE enriches at nuclear pores labeled by NUP62 and imaged by STED super-resolution microscopy (30nm xy-pixel resolution). Shown are linearized regions from cross-section images of the nuclear envelope (scale bar below), comparing PE(18:1/Y_16_) to PC(18:1/Y_16_) and pan-lipid metabolic labeling using Y_16_, which exhibits more uniform signal along the nuclear envelope than either phospholipid species. **L)** Particle average images of lipid localization at nuclear pores from the basal (flattest) plane of the nuclear envelope surface. Shown are the mean intensity patches from 390×390nm areas generated by particle averaging of NUP62 maxima, for both lipid (*top*) and NUP62 (*lower;* average of N>2000 pores from N≥6 nuclei from N≥2 biological replicates per condition). PE enriches around the nuclear pore at the expense of the surrounding membrane, and especially closer to the center of the pore, a trend that is less pronounced for PC(18:1/Y_16_) or palmitic acid (Y_16_) See also **Extended Data Figure 7**. **M)** Quantification of lipid signal in the inner donut (60nm radius excluding pore center) from particle averaged images as in **L**, shown as the percentage of pixels with high lipid signal compared to the surrounding nuclear envelope (≥160nm from pore center). PE exhibits a ∼6-30% enrichment compared to PC and pan-lipid labeling by Y_16_ (**p<0.01, ***p<0.001 by one-way ANOVA with multiple comparisons).

Based on our findings, we predicted that the MICOS and nuclear pore complexes interact with PE lipids within their highly curved resident membranes domains via the core transmembrane proteins MIC60 and NDC1. To test this hypothesis, we first assessed whether MIC60 and NDC1 are direct PE interactors. From lipid pulldowns across time, we found that PE rapidly and consistently associates with both proteins, surpassing our quantification thresholds at all timepoints (**Figure 4E-F**). Very little of the PE(18:1/Y_16_) probe has reached either organelle at 1 minute, as assessed by imaging, and most other mitochondrial or ER proteins do not exhibit a similar level of early enrichment (**Extended Data Figure 6A-F**). This indicates that PE has a high preference for associating with MIC60 and NDC1. Moreover, both interactions were resistant to detergent treatment prior to pulldown elution (**Figure 4G-H, Supplemental Table 5, Extended Data Figure 6G-H**), while the same trends were not observed for peripheral protein components of each complex, such as MIC25^98^ or ALADIN^99^. MIC60 and NDC1 are thus likely direct interactors that selectively associate with PE at the expense of other lipids and proteins.

Next, we sought to visualize the lipid-protein interactions in their native membranes by high-resolution imaging. For the MICOS complex, we used spinning disc confocal microscopy at 150X to map PE(18:1/Y_16_) co-localization with inner versus outer mitochondrial membrane proteins. PE is observable at cristae-like subdomains within the filamentous mitochondrial structure, co-localizing with both MIC60 and MIC25 more than the outer membrane proteins TOMM20 and MAVS at all timepoints (**Figure 4I-J, Extended Data Figure 7A-B**). This is especially pronounced at 20 minutes, when the MIB complex exhibits the highest overall enrichment by PE (**Figure 4E, Extended Data Figure 6C**). For the nuclear pore complex, we used STED super-resolution imaging of PE(18:1/Y_16_) at the nuclear envelope, targeting nuclear pores via antibodies against endogenous NUP62 (**Extended Data Figure 7C-E**). PE exhibited heterogenous signal along cross-sections of the nuclear envelope rim, often enriching at NUP62 puncta (**Figure 4K**). We also compared to PC(18:1/Y_16_) and bifunctional palmitic acid (Y_16_), which represents a pan-lipid membrane label as it is rapidly metabolized into numerous lipid species. PC signal was also heterogenous, although the brightest regions appeared more often in nuclear envelope regions that were not co-localized with NUP62. The pan-lipid Y_16_ exhibited more uniform distribution compared to both phospholipid species, indicating that we are resolving lipid-specific nanoscale localization at the nuclear envelope. Using nuclear surface images, we performed particle averaging of NUP62 maxima to resolve nuclear pores in the lipid membrane and quantify lipid signal in the pore vicinity (**Figure 4L-M, Extended Data Figure 7H-J**). We find that PE is enriched at the nuclear pore and depleted from the surrounding nuclear envelope, especially at regions corresponding to the apex of the pore membrane (<60nm), which is the expected location of NDC1 (**Extended Data Figure 7H**). In contrast, PC(18:1/Y_16_) and especially Y_16_ are more similarly distributed across the nuclear membrane, with similar lipid intensities observed in the pore vicinity compared to the surrounding nuclear surface. We thus provide direct in-cell evidence for selective PE species enrichment at curved membrane subdomains which are critical to the structural integrity of mitochondrial cristae and the nuclear envelope.

## PE stabilizes MIC60, NDC1 at curvature

Given the prevalence of the PE-MIC60 and PE-NDC1 interactions within the complex cellular environment, and the known capacity for PE to enable membrane deformations, we hypothesized that PE(18:1/Y_16_) may act to position each protein within its highly curved membrane domain. As both NDC1 and MIC60 are essential for assembly of their respective complexes and thus the formation of the nuclear pore^85,100–102^ and cristae junctions^93,98,103,104^, genetic perturbation experiments would be difficult to interpret. We instead turned to coarse-grained molecular dynamics (MD) simulations of model membranes to further characterize the lipid-protein interactions and their influence on transmembrane protein positioning within curved membranes (**Figure 5**).

**Figure 5.**
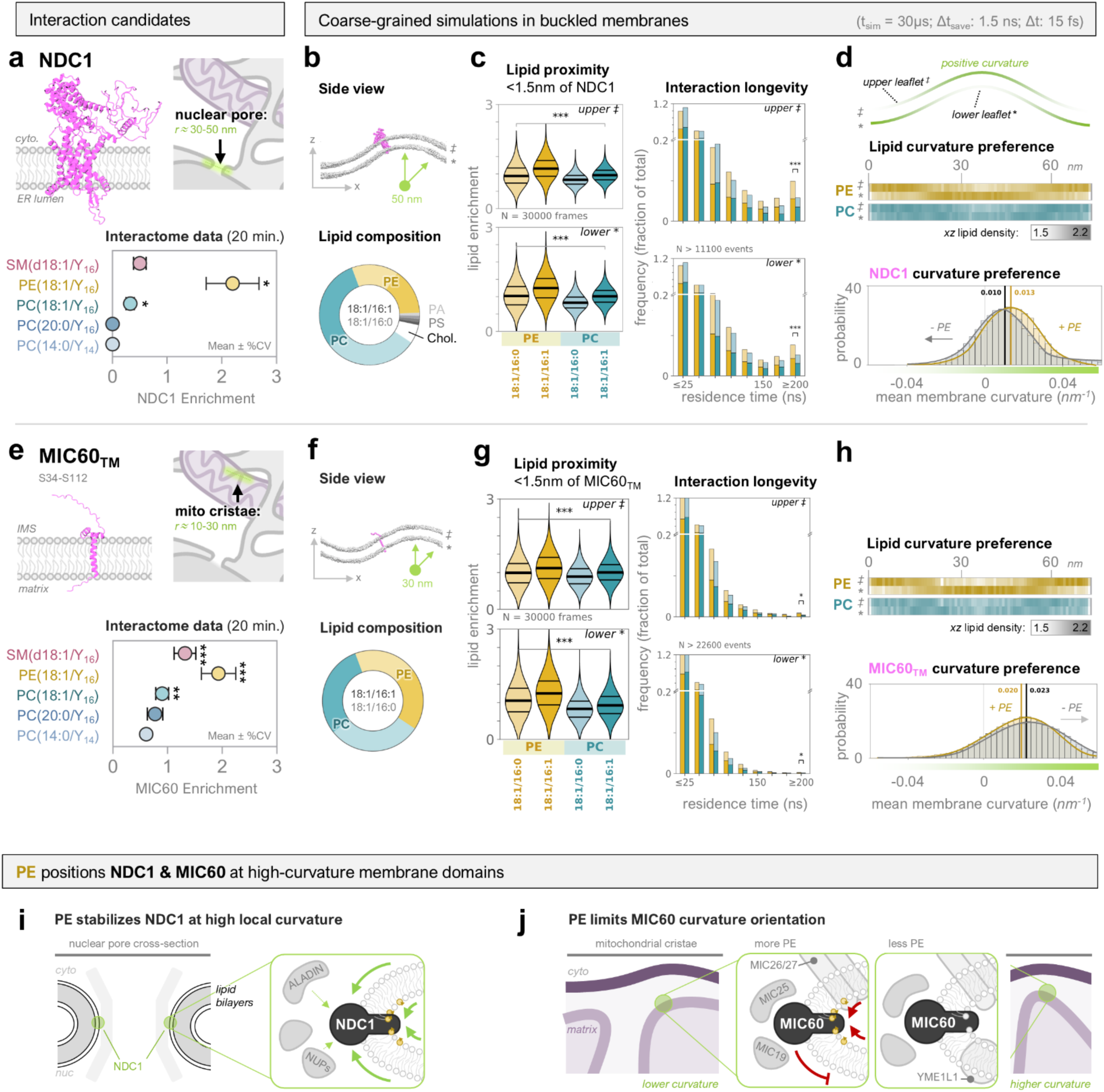
PE interactions stabilize MIC60 and NPC1 at curved membrane nanodomains. **A)** Structure of full-length NDC1 used in MD simulations (*left*, derived from PDB ID: 7R5J^92^) and cartoon of organelle localization indicating the mean radius of curvature at resident membrane domains (*right*). The mean protein enrichment from the lipid pulldowns is *below* (as in Figure 2; *p<0.05, **p<0.01, ***p<0.001 Benjamini-Hochberg corrected). **B)** *Top,* Side view of the simulated buckled membrane (r=50nm, representative single timeframe of *xz*-axis; lipid headgroups = grey, proteins = magenta). *Bottom,* Donut plot of lipid composition. Each lipid class includes equimolar ratios of palmitoleic (16:0/18:1) and di-oleic (18:1/18:1) derivatives; PC (teal) and PE (yellow) are highlighted in bright colors matching the interactome datasets. For NDC1 simulations, the membrane is “nuclear envelope (NE)/ER-like”, with PE species comprising 30% and PC species 60% of the total lipids and r=50nm. Simulations with other lipid compositions are included in **Extended Data Figure 8**. Simulations were 30µs (Δt = 15 fs), and each replicate started with the proteins initially positioned at different curvatures (positive, neutral, negative; N=3). **C)** Quantification of NDC1 interactions with PE (yellow) and PC (teal) lipids at the upper (‡, *top*) and lower (*, *bottom*) leaflets of the bilayer. Lipid proximity is the residence fraction of each lipid species <1.5nm from the protein in each frame (N = 30,000), shown as a violin plot (thick line = median; ***p<0.0001 by Brown-Forsythe multiple comparisons test). Interaction longevity is reported as a histogram of the fraction of lipids remaining <0.6nm from the protein for >1.5ns (bin size = 25ns, lipid binding events N>11100). PE outcompetes PC for proximity throughout the simulations, especially for 18:1/16:1 species and at the lower leaflet, and PE interactions are longer-lived (≥75 ns) at both leaflets (***p<0.0001 by two-proportion z-test). **D)** Positioning of lipids (heatmaps, *top*) and proteins (histograms, *bottom*) as a function of membrane curvature (simulation structure plotted at top, *x*-axis to nm scale with heatmaps, green gradient = positive (green) to negative (white) curvature). For the heatmaps, the maximum lipid density at each x coordinate (averaged across *y* and time, N=30,000 frames) is shown for both the upper (‡) and lower (*) leaflets of the bilayer, with PE(18:1/18:1) in yellows and PC(18:1/18:1) in teals plotted on the same scale (*grey* key below). In every simulation, PE enriches at the lower leaflet of positive curvature. For the histograms, the frequency distribution of protein curvature residence (*nm*^-1^, *x*-axis) at every timepoint is shown (yellow; N=30,000; bin size = 0.003 *nm^-1^*, median is labeled). NDC1 exhibits preference for positive curvature in the presence of PE. Loss of PE shifts curvature preference by 3%, shown as a grey overlay of the corresponding histogram from simulations containing 0% PE (“-PE”, arrow indicates direction of shift). **E)** Structure of MIC60_TM_ (residues S34-S112) used in MD simulations, cartoon of organelle localization, and interactome data, shown as for NDC1 in **A.** The transmembrane domain (I46-Y64) plus flanking residues was used for MIC60 due to lack of high-confidence human structures, predicted by AlphaFold3 (see **Extended Data Figure 8** and **Methods**). **F)** Side view and “inner mitochondrial membrane (IMM)-like” lipid composition of simulated buckled membranes (r=30nm), shown as in **B.** Relative PE lipid content is increased to 40% to reflect the greater proportion of PE and lower amounts of other phospholipids in the IMM; see **Extended Data Figure 8** for simulations of other lipid compositions including with cardiolipin **G)** Quantification of the proximity (*left*) and longevity (*right*) of lipid interactions with MIC60_TM_, plotted for the upper (‡, *top*) and lower (*, *bottom*) leaflets as in **C** (lipid binding events N>22600). PE outcompetes PC for proximity throughout the simulations, especially for 18:1/16:1 species and at the lower leaflet. PE interactions are longer-lived (≥75 ns) at both leaflets (*p< 0.05 by two-proportion z-test). **H)** Lipid (heatmaps) and protein (histogram) curvature preferences, shown as in **D**. MIC60_TM_ exhibits preference for positive curvature in the presence of PE, and a further 3% increase in curvature preference upon removal of PE (“-PE” yellow histogram, arrow indicates direction of shift). **I-J)** Models for the topology and function of PE interactions with NDC1 and MIC60 at the nuclear pore and mitochondrial cristae, respectively, based on the biochemical, imaging, and MD simulation evidence from this study and placed in the larger biological context.

Specifically, we simulated full-length NDC1 and the transmembrane region of MIC60 (MIC60_TM_) in buckled (*r* = 50 and 30 nm, respectively) membranes comprised of lipid species reflecting simplified organelle membrane environments (**Figure 5A, E**). This included palmitic (16:0), palmitoleic (16:1), and oleic (18:1) acid derivatives for each structural lipid class (**Figure 5B, F**), as we previously showed that the bifunctional lipid behaves more similarly to acyl chains with one unsaturation (e.g., Y_16_ is more similar to 16:1)^14^. We also included TMEM147, an ER sheet-resident protein that was selectively enriched by PC species at the expense of PE in our pulldown datasets and has no known curvature preference (**Extended Data Figure 9A-D**). For every protein, we ran simulations in three lipid environments: 1) with or 2) without PE, and 3) with additional PI (nuclear envelope-like) or cardiolipin (mitochondria-like) (see **Methods** and **Extended Data Figure 8**). In addition, we tested three starting positions for each protein: positive, neutral, or negative curvature.

Quantification of lipid-protein interactions showed that PE lipid species out-compete PCs for interactions with NDC1 and MIC60_TM_, especially at the lower leaflet of the membrane bilayer (**Figure 5C, G**). PE-protein interactions were also more stable, persisting in longer-lived contacts with both proteins for hundreds of nanoseconds, especially for NDC1. We note that entropic effects are introduced in complex membrane mixtures. Therefore, even slight enrichment of a certain lipid around a protein is indicative of lipid preference. In addition, NDC1 and MIC60_TM_ primarily localized to positive curvature in our simulations (**Figure 5D-H**). In contrast, TMEM147 did not exhibit curvature preference, and interacted with PE and PC species at similar levels (**Extended Data Figure 9F-G**). PE also accumulated at the inner leaflet of the buckled membrane (**Figure 5G-H**). When the proteins are embedded in membrane environments containing highly charged lipids with strong conical shapes, such as PI for NDC1 and TMEM147 or cardiolipin for MIC60_TM_, there is a preference for those lipid types, which are not included in the interactome data (**Extended Data Figure 8**). Altogether, these findings confirmed our pulldown data and suggested that NDC1 and MIC60 experience either protein-intrinsic or lipid-directed effects on curvature preference, specifically at the inner leaflet as predicted for a conical-shaped lipid.

When we removed PE from the simulations, we observed a shift in the curvature distribution of each protein. For NDC1, which displays the longest-lived interactions with PE, the absence of PE led to a shift in the distribution toward neutral curvature. For MIC60, which has a lower fraction of long-lived PE interactions, the absence of PE shifted the distribution toward higher curvature values (**Figure 5G-H)**. We also observed an effect for TMEM147, which shifts into negative curvature regions in the absence of PE, overall suggesting that PE differentially influences the preference of transmembrane proteins for curved membrane domains. Moreover, when we flattened the simulated membrane to remove the influence of membrane curvature on lipid-protein interactions, we still observe NDC1 and MIC60_TM_ preferentially interacting with PE, while TMEM147 displayed preferential interaction with PC (**Extended Data Figure 9H-J**). Therefore, PE-NDC1 and PE-MIC60 interactions are specific rather than a product of proximity driven by intrinsic preferences for membrane curvature. Altogether, our findings lead us to propose that interactions with PE differentially regulate NDC1 and MIC60 via both the biophysical preference of PE for curved membranes and direct, specific interactions with both proteins. PE stabilizes NDC1 within high curvature membrane nanodomains, whereas PE limits the preference of MIC60 for increasing curvature (r≤30nm). These dual mechanisms may serve to control the local protein densities needed for functional membrane architecture.

## Discussion

Membrane function is primarily attributed to proteins, yet all proteins coevolved with lipids, and the cell expends significant resources producing, regulating, and organizing thousands of unique lipid species. Mechanistic studies of lipid cell biology have remained scarce, as methodological development has for decades been focused on the biomolecules of the central dogma (nucleic acids and proteins). Here, we present an experimental blueprint for systematically defining the protein interactomes of individual membrane lipids. Using bifunctional phospholipid probes for photoaffinity capture proteomics in combination with super-resolution lipid imaging and molecular dynamics simulations, we identify and functionally characterize selective lipid-protein interactions in organelle membranes.

Our study leads us to two general conclusions for membrane biology: first, nearly 70% of the 668 top interaction candidates that we identified were specific to a single lipid species, uncovering a remarkable selectivity that extends across protein pathways, complexes, and changing organelle localizations (**Figures 2-3**); and second, common lipid species engage in more selective protein interactions than rare ones—findings that mark the regulation of subcellular lipidome complexity as critical for building specialized membrane co-assemblies. Achieving the molecular precision captured by our interactome datasets required accounting for the confounding features of lipid biology, including metabolic interconversion, inter-organelle lipid transport, regulation of protein abundance, and widely varying lipid-protein affinities. Our LP-AP framework is built on four core experimental considerations (**Figure 1**): i) Monitoring individual phospholipid species using pulse-chase experiments, which provides key spatiotemporal resolution for tracking the development of lipid transport, metabolic turnover, and protein interactions; ii) comprehensive proteomic quantitation of enriched proteins alongside whole-cell proteome changes and background contaminants (FLUB), which accounts for lipid-induced protein changes and promiscuous membrane biochemistry; iii) retention of direct interactors together with associated peripheral proteins, which enhances information content and informs on the topology of lipid-protein complexes; and iv) comparative analysis of multiple lipid species in parallel, which all control for each other. Through multimodal experiments that comprehensively address these issues, we advance beyond earlier studies, including our own^41^, to generate informed hypotheses on lipid function across the many cellular membrane compartments and their rich membrane proteomes.

In two case studies, we followed up on interactions between PE and the core ancestral integral membrane subunits of membrane organizing complexes at mitochondrial cristae (MIC60; MICOS) and the nuclear pore (NDC1; NPC), which are both conserved across nearly all eukaryotes^83,84,86,100^ (**Figures 4-5**). We find that PE co-assembles with NDC1 and MIC60 at their curved resident membranes *in vivo* and directly interacts with both proteins *in vivo* (**Figure 4**) and *in silico* (**Figure 5**). Both super-resolution imaging and MD simulations show that PE increases in density at the membrane segments of highest curvature and stabilizes the localization of both proteins at those domains.

Interestingly, removing PE from the simulations has opposite effects: NDC1 loses while MIC60_TM_ gains preference for increasingly positive curvature. This points to protein-intrinsic determinants of the functional output of PE interactions, a finding that represents the different biology of each protein. For example, although NDC1 prefers high-curvature membranes *in vitro*^101^, it does not drive membrane curvature itself and requires other curvature-inducing proteins to properly localize *in vivo*^105^. Conversely, MIC60 autonomously curves membranes into thin *r*=5-10nm tubules *in vitro*^93,103,106^, and is the only transmembrane MICOS component that can assemble independently of cardiolipin^107^, an exclusively mitochondrial lipid with intrinsic curvature exceeding that of PE^88,108,109^ (**Extended Data Figure 8**). In addition, nuclear pores are largely stable following assembly (*r*=30-50nm)^97,110–112^, while cristae are frequently remodeled within a much greater curvature range (*r*=10-40nm)^93,94,98^. NDC1 targeting to and subsequent anchoring of newly-made NPCs at *de novo* membrane pores is an essential step in both interphase and post-mitotic nuclear pore assembly^85,97,100–102,113–115^, and one study implicated PE in this process^116^. Alternatively, aberrantly high PE levels “loosen” cristae architecture and disrupt overall mitochondrial function^109,117–119^, likely mediated by the PE-regulated protease YME1L1^64,65,120,121^. Therefore, we propose a model whereby PE stabilizes NDC1 at high local curvature during anchoring of the NPC anchoring to the nuclear envelope, while limiting the extent to which MIC60 can curve membranes during cristae remodeling (**Figure 5I-J**). Our in-cell data indirectly supports these dual mechanisms, showing that PE has multiple direct interactors within the MICOS and co-localizes with both mitochondrial membranes, while NDC1 is the only direct interactor and the only transmembrane NPC protein detected (**Figure 4**). The PE-MIC60 and PE-NDC1 co-assemblies are best understood as a combination of the selective nano-environment of a curved membrane, which increases local PE density, and discrete lipid-binding events, which regulate the positioning of both proteins via longer-lived interactions. Taken together, our findings expand the fundamental architecture of both complexes to include PE lipids around the transmembrane domains of their protein cores, particularly at the inner leaflet of the curved bilayer. This begins to address several long-standing questions with regard to the relationship between localized lipid metabolism and organelle shape, particularly for the nuclear envelope^122,123^.

Our experimental framework is generalizable to other lipid species, MS acquisition methods, biochemistry (e.g., organelle isolation), computational analysis pipelines, and biological contexts. Moving forward, we perceive three major bottlenecks: i) the chemical synthesis of more bifunctional lipid species for different biological contexts, which is time-consuming; ii) navigating biological processes that occur at longer timescales, when metabolic breakdown of the chosen probes will become significant; and especially iii) identifying the actual crosslinking sites of lipid-protein conjugates, all of which necessitate continuous methodological development.

In summary, our work translates the chemical diversity of membrane lipids into species-resolved maps of supramolecular membrane complexes, with functions exerted by both protein and lipid components. We expect that our publicly available datasets will serve as starting points for many cell biological investigations but also to computationally map protein-intrinsic features that exhibit selectivity for different lipid species, such as transmembrane domain lengths, widths, and flanking domains^7,124^. The underlying methodology is suitable for both characterizing the protein components of lipid sorting compartments, as we showed for SM interactions at non-canonical vesicles, and the direct regulation of membrane proteins by individual lipid species, as demonstrated for the MICOS (MIC60) and nuclear pore (NDC1) complexes by PE. Altogether, our approach enables effective investigations of the molecular functions of individual lipid species in cell biology.

## Supporting information

Supplemental Table 5 Harsh Wash

Supplemental Table 6 WCL Data

Supplemental Table 1 FLUB

Supplemental Table 3 Timecourse

Supplemental Table 4 Annotations

Supplemental Table 2 5 Lipids

## Acknowledgements

We are grateful to A. Knaust (Shevchenko group) for excellent MS quality control and D. LaJoie (Human Technopole) for valuable advice regarding nuclear pores; the Light Microscopy (B. Schroth-Diez and J. Peychl), Workshop (F. Elsner), and Genome Engineering (I. Reichardt-Gómez and M. Sarov) Facilities at MPI-CBG (Dresden) for imaging resources, photoreactor engineering, and generation of stable cell lines, respectively; and the Max Planck Computing and Data Facility for computing resources for MD simulations. We thank M. Marass for providing editorial advice during the publication process. We acknowledge funding from the European Research Council to AN (Horizon 2020 GA758334 ASYMMEM and AURORA) and AvA (ORGANELLOIDS 101117619); the Deutsche Forschungsgemeinschaft to AN, AH, and AS (TRR83 consortium) and AvA (SPP2191); the Paul G. Allen Family Foundation to AN and AH.; the Heineman Foundation to AvA (Minerva); an ELBE postdoctoral fellowship to SL; the “Hessen Horizon Marie Skłodowska-Curie-Stipendium” program to KPR; and the Max Planck Society for support of KCC, KPR, KB, HML, SL, AS, GH, AvA, and AN.

## Author Contributions

KCC designed the LP-AP pipeline with AN and experimental follow-ups with AN and AvA. KCC established the LP-AP workflow; KB synthesized all bifunctional lipid probes; KCC collected, acquired and analyzed MS samples; KCC interpreted MS data with AS and SL; KCC and AN evaluated biological findings; KCC acquired and analyzed spinning disc confocal images; KCC and HML acquired and analyzed STED images with AH; KPR designed, ran, and analyzed MD simulations with GH. KCC and AN wrote the manuscript; KCC prepared figures, including cartoons; all authors read and commented on the manuscript.

## Conflict of Interest

The authors declare no competing interests.

## Data Availability

The complete, annotated LP-AP interactomes, FLUB assignments, harsh detergent wash analysis, and whole-cell proteomes are included in Supplemental Tables 1-6 as detailed Excel dataframes, serving as comprehensive resources for lipid studies. All MS raw data has been deposited with to ProteomeXchange via PRIDE (EMBL-EBI); during the review process this can be accessed via username and password upon request to the corresponding authors. Coarse-grained MD trajectories for each replicate and initial configuration will be deposited in the Zenodo entry XXX [https://zenodo.org/records/XXX]; during the review process this can be accessed via OwnCloud upon request. For further information, resources, or reagents, contact the corresponding authors.

## Extended Data

**Extended Data Figure 1.**
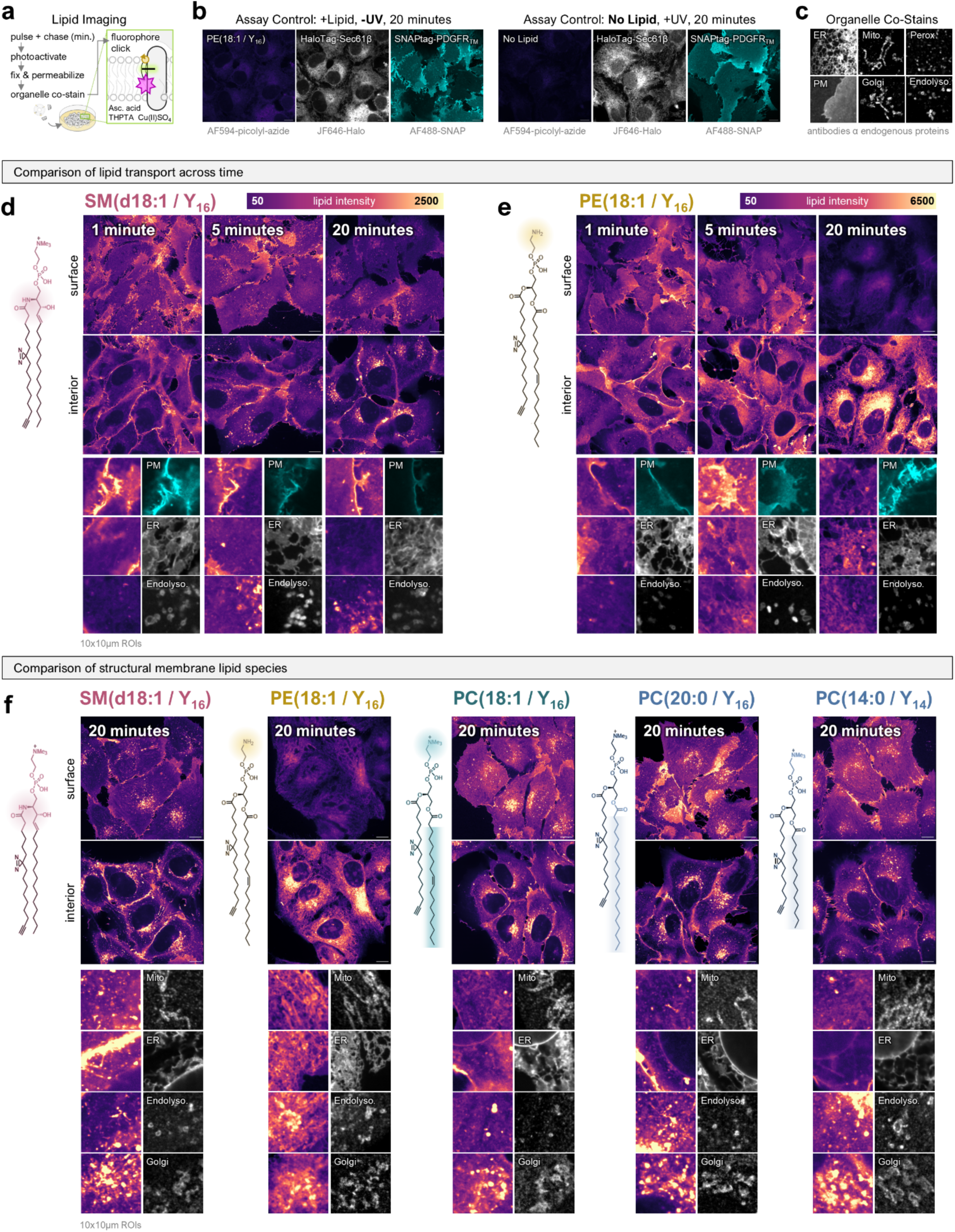
Time-resolved organelle localization of membrane phospholipids. **A)** Lipid imaging workflow schematic. At the desired timepoint after bifunctional phospholipid addition (e.g., 5 or 20 minutes), the diazirines are photoactivated with a 1 sec. UV pulse using high-power LEDs to anchor the lipid probes to the cellular protein network. Samples are then immediately chemically fixed, permeabilized, and immunostained for organelle proteins. Finally, lipids are visualized via addition of AlexaFluor594-picolyl-azide by click chemistry. See also Figure 1, **Methods**, and Iglesias-Artola *et al.* 2025^14^. **B)** Imaging controls include a -UV (no photoactivation) condition to monitor probe crosslinking efficiency and a no lipid (NL) condition that is also UV irradiated, co-stained, and copper clicked to monitor background signal in the lipid channel. The respective ER (HaloTag-Sec61β; labelled with JF646-Halo) and plasma membrane (PM; SNAPtag-PDGFR_TM_; labelled with AF488-SNAP) co-stains are shown for representative images of U2OS cells (100×100µm field of view). **C)** Example 10×10µm regions of organelle co-stains from this study. Except for the ER and PM (labeled by stable transgene expression), organelle population co-stains used antibodies against endogenous organelle proteins: mitochondria were labeled with a mix of TOMM20 and MAVS, peroxisomes with PEX14, Golgi with GM130, and endolysosomes with a mix of RAB5, RAB7, and Lamp1. See also **Methods.** **D)** Representative images of SM(d18:1/Y_16_) in U2OS cells at 1, 5, and 20 minutes post-lipid pulse (0, 1.5, and 16.5 minute chase times) using 150X spinning disc confocal imaging. The bifunctional lipid probe structure is shown at *left*. All images are acquired as 4-channel, 0.3µm z-stacks; a PM surface image and mid-cell interior image from the same z-stack is shown for each timepoint. Organelle co-stains alongside the corresponding lipid channel are shown as regions of interest (ROIs) below. For direct comparison across timepoints, all images are shown on the same intensity scale (key at *top*). Scale bars and ROIs are 10µm in width. **E)** Representative images of PE(18:1/Y_16_) at 1, 5, and 20 minutes post-lipid pulse, displayed as in D. **F)** Comparison of the organelle localization of five structural membrane phospholipids at 20 minutes post-lipid pulse in U2OS cells. The structure of each bifunctional lipid probe is shown at *left* of the corresponding images, and representative ROIs for organelle co-stains *below*.

**Extended Data Figure 2.**
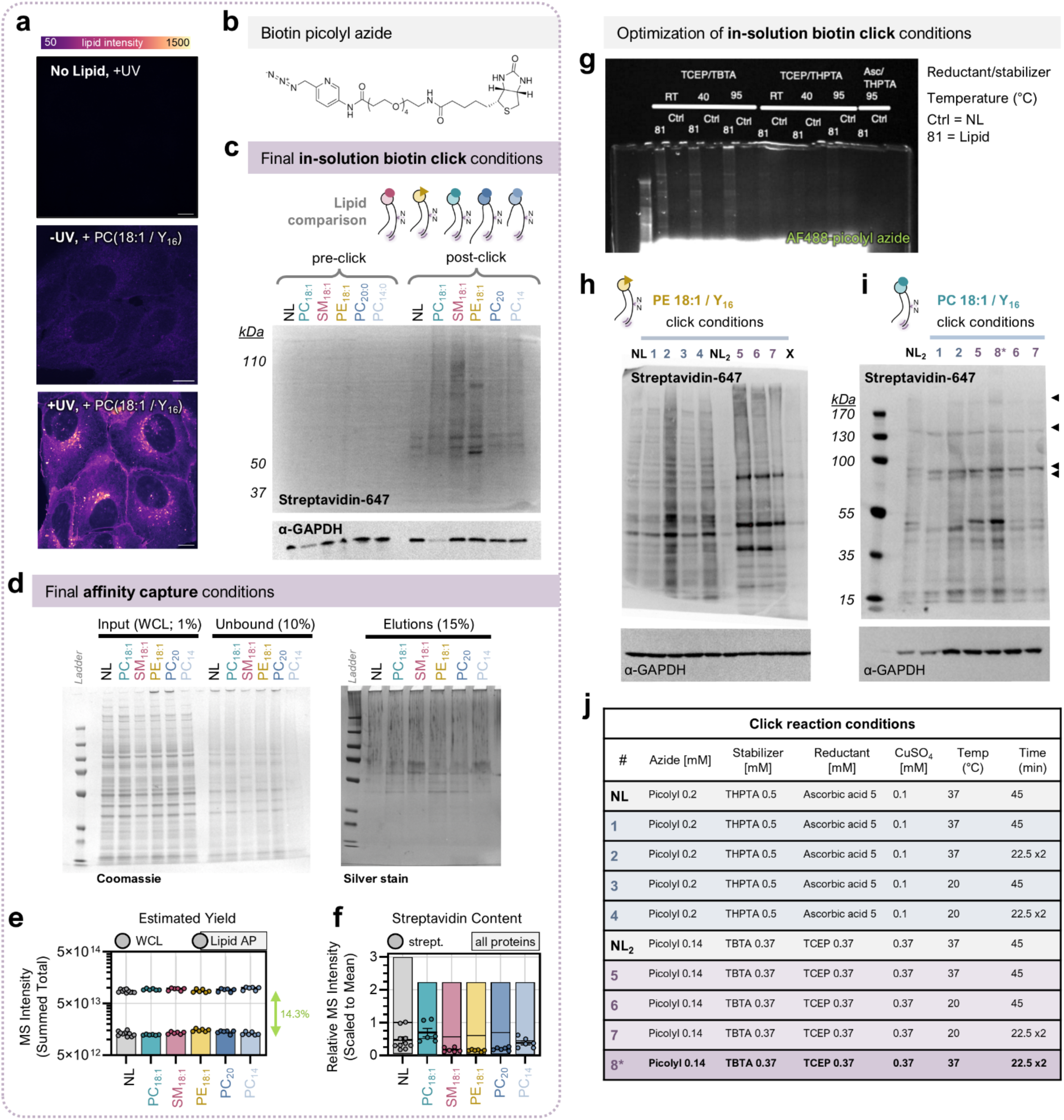
Optimization of the lipid-protein affinity pulldown (LP-AP) biochemistry. **A)** Example lipid channels from PC(18:1/Y_16_) imaging experiments used to inform control selection for LP-AP. Note that the -UV condition was not used as a “control”, as this condition would capture lipid-protein interactions whose stability does not require diazirine crosslinking. All images are shown on the same intensity scale for direct comparison of signal. Scale bars are 10µm. **B)** Structure of the biotin picolyl azide used for copper-mediated in-solution click reactions in this study (Jena, CLK-1167). The picolyl azide group enhances reaction specificity and efficiency^125^. **C)** Western blot of biotin content in LP-AP samples before (*left*) and after (*right*) in-solution biotin click, labelled via Streptavidin-647 fluorescent conjugate. No lipid (NL) controls are compared to the five structural phospholipids analyzed in this study: PC(18:1/Y_16_), PE(18:1/Y_16_), SM(d18:1/Y_16_), PC(20:0/Y_16_), PC(14:0/Y_16_) from *left to right* (shorthand notation is used in the figure). NL controls always have click background; the lipid-specific content varies by lipid species. GAPDH is shown *below* as a loading control from the same blot. **D)** Protein SDS-PAGE gels indicating the assay efficiency of the final affinity capture conditions used in this study, comparing the NL control to all five lipid species. *Left*, Coomassie stain of total protein content in the starting input lysate (*left*) and unbound sample (*right*) following biotin-streptavidin isolation. *Right*, Silver stain of total protein content following elution. Eluates are difficult to visualize by SDS-PAGE due to distribution of lipid-protein conjugates across molecular weights (faint “streaks” in sample lanes); the amount of material was more than sufficient for MS analysis. **E)** MS quantification of sample content, comparing whole-cell lysates (WCL; dots) to LP-AP eluates (dots + bars), which were approximately an order of magnitude lower in abundance. Data is shown as a scatter plot of the summed MS intensity (*y*-axis) per lipid condition (*x-*axis), each dot is a replicate (N=6). Average “yield”, or comparative sample abundance, is ∼14.3% for LP-AP samples from their input WCLs. **F)** MS quantification of the relative streptavidin content per LP-AP sample, shown as a box plot where every dot is a replicate (N=6 for lipids and N=10 for NL; bold line at median, error bars are standard deviation, min-max are 10-90 percentiles). Streptavidin should enter the eluate due to the use of a low-pH elution from the beads; streptavidin content compared to endogenous proteins is minimal (<<1). **G-J)** Comparison of in-solution click conditions for lipid-protein conjugates following our lysis conditions. **G)** In-gel fluorescence visualization of fluorophore-click following different reaction conditions: TCEP/TBTA, TCEP/THPTA, or Ascorbic acid/THPTA (used for lipid imaging) at either room temperature, 40°C, or 95°C. Ctrl = no lipid (NL) control; 81 = bifunctional PC lipid. **H)** Western blot of biotin in NL control versus PE(18:1/Y_16_) or **I)** PC(18:1/Y_16_) lipid click reactions, comparing different click conditions outlined in the table in **J.** Ultimately, TCEP/TBTA at physiological temperatures performed best, with lower NL background observed and consistently higher biotin click efficiency. Condition 8* was chosen as the final click conditions for all experiments (such as in **C**). Black triangles indicate molecular weights of proteins that likely correspond to endogenous carboxylases (ACC, PC, 3-MCC, PCC), which can be biotinylated.

**Extended Data Figure 3.**
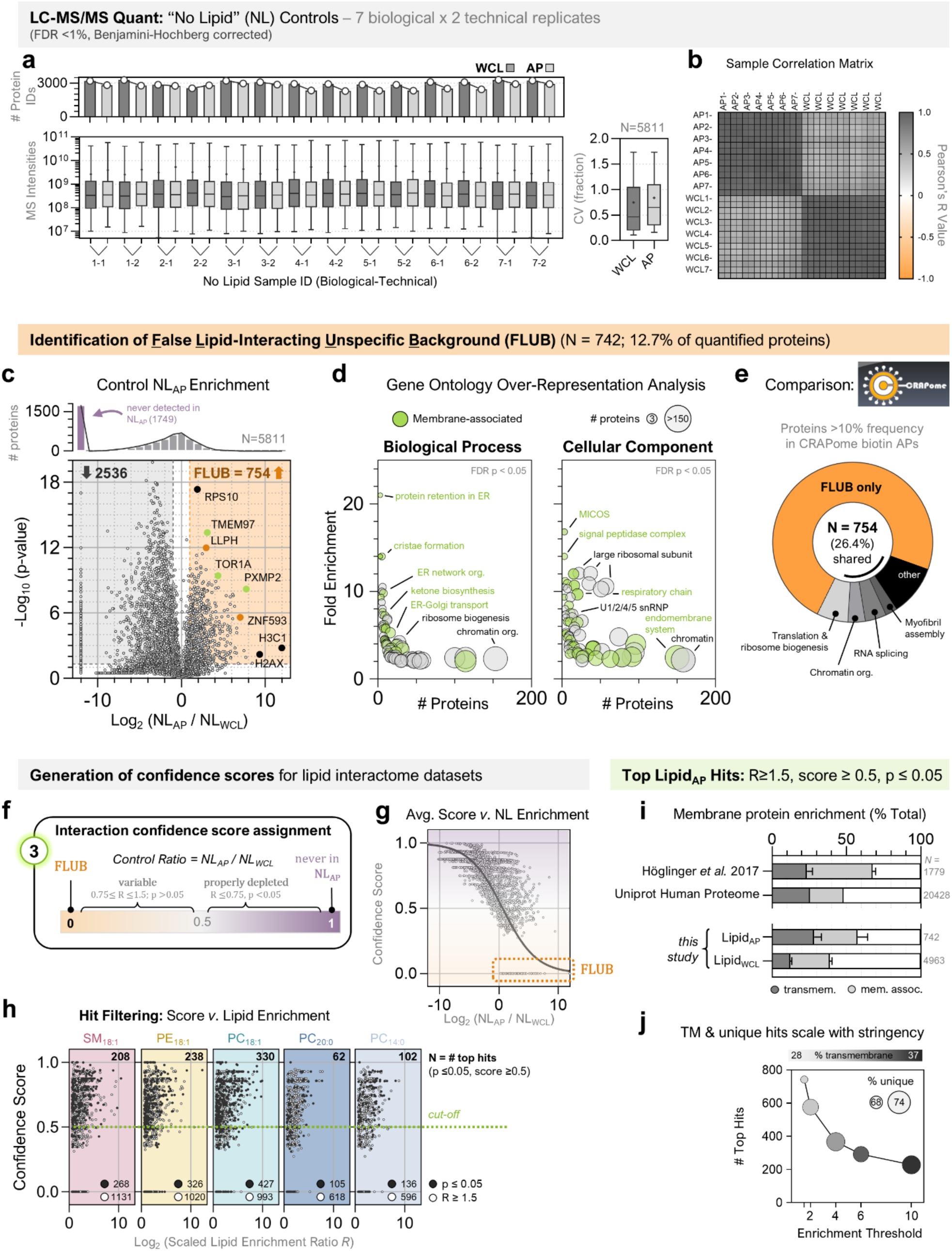
Characterization of <u>F</u>alse <u>L</u>ipid-Interacting <u>U</u>nspecific <u>B</u>ackground (FLUB) and use for scoring of interaction confidence. **A)** Summary of LC-MS/MS datasets resulting from no lipid (NL) controls in our study (N=14 total, 7 biological x2 technical replicates). *Top*, the number of high-confidence protein IDs in whole-cell lysate (WCL; dark grey) and AP (light grey) samples; *bottom*, the distribution of MS intensities for quantified proteins (Benjamini-Hochberg corrected FDR < 1%, minimum 2 unique peptides per protein, see **Methods**); *right*, the coefficients of variation (CV) for all proteins following averaging of replicates. Data is shown as box-and-whisker plots with line at median, + at mean, whiskers 2.5-97.5%. Note that the CVs are higher for NL APs than for lipid APs (see **Extended Data Figure 4**), in-line with a background control. **B)** Sample correlation matrix of all WCL and AP samples from NL controls (technical replicates combined), with positive correlation (Pearson’s R value) shown in grey and anti-correlation in orange. NL_AP_ samples are more similar to each other than NL_WCL_ samples. **C)** Volcano plot of protein enrichment in NL_AP_ versus NL_WCL_ samples (N=14 replicates averaged; N=5811 proteins), with the enrichment ratio as *x* and the p-value (-Log_10_ transformed, Benjamini-Hochberg corrected) as *y.* Proteins that are significantly enriched (R>1.5, p<0.05) in NL control APs are in the shaded orange region at *right*; these are named FLUB (N=754 proteins), with several examples labeled (black = also in CRAPome, orange = FLUB only, lime = known membrane association). *Top*, histogram showing the number of proteins across the *x*-axis, with many capped at far *left* indicating proteins that were never detected in NL_AP_ (N=1749) but were present in either WCL or Lipid_AP_ samples. D) FLUB proteins were assessed for over-represented gene ontology terms using PAN-GO^57^ biological process (*left*) or cellular component (*right*). Data is shown as multi-variable plots whereby the number of FLUB proteins assigned to each term is on the *x*-axis, the fold-enrichment compared to the human proteome is on the *y*-axis, the node color indicates a known membrane-associated function (lime green), and the node size again indicates the number of FLUB proteins assigned to the category. Some prevalent examples are labeled; all terms were already filtered by FDR-corrected p-value. We note that that some subunits of the MICOS complex (3 proteins in total) are assigned to FLUB, in line with previous reports of “stickiness”; these do not include the high-confidence PE interactor MIC60. **E)** FLUB proteins were compared to common contaminants in the CRAPome repository for protein-protein interaction studies. 199 (26.4%) of the 754 FLUB proteins were reported with ≥10% frequency in proximity biotinylation control experiments in human cells (black/grey donut segments); these are further categorized into major biological categories such as translation and chromatin organization, to show shared characteristics of FLUB and CRAPome proteins. See also **Supplemental Table 1.** **F)** Schematic showing the use of NL datasets for scoring interaction confidence, whereby FLUB proteins receive score=0 and proteins never detected in NL_AP_ (N=1749) receive score=1; the remaining proteins from the NL datasets are scaled from 0 to 1 based on significance and enrichment ratio. The number 3 represents the third dataset used for LP-AP analysis, alongside Lipid_AP_ and Lipid_WCL_ datasets (see Figure 1). **G)** Final LP-AP confidence scores applied to proteins in LP-AP datasets, shown here for the NL control (*x* = enrichment ratio, *y* = score) as a scatter plot. Proteins that are less abundant in NL_AP_ receive higher scores, as these are proteins that are properly depleted. Note that the final score is the average score from all individual NL replicates (see **Methods**). **H)** Scatter plots showing the application of confidence scores to the Lipid_AP_ datasets (as in Figure 2), with the scaled enrichment ratio on the *x*-axis (positive values only) and assigned score on *y.* Each lipid species is indicated at top and color-coded as in all figures (shorthand lipid notation is used here). A cut-off score of 0.5 was applied for selection of top hits. For each lipid, the final number of top hits is indicated at the top of each graph (e.g., 208 for SM), with the original number of enriched proteins (R≥1.5) indicated in white dots and significantly enriched proteins (R≥1.5, p≤0.05) prior to scoring indicated as black dots. **I)** Following confidence scoring, top hits from Lipid_AP_ datasets (N=742) were annotated by membrane association (transmembrane=dark grey, associated/peripheral=light grey) and compared to all proteins from Lipid_WCL_ in this study, to the total human proteome (Uniprot, N=20428), or to reported hits from Höglinger *et al.* 2017^41^ (N=1779). Data is shown as a stacked bar plot (error bars = standard deviation). **J)** Analysis of the relationship between transmembrane domains, lipid-specific interactions (unique), and enrichment ratios for top hits across the five lipids in our study (*y*-axis), shown as a multi-variable plot. Increasing enrichment thresholds (e.g., >4-fold enrichment compared to NL controls, *x*-axis) result in a higher proportion of transmembrane proteins (color scale) and proteins only detected as interactors for a single lipid species (node size).

**Extended Data Figure 4.**
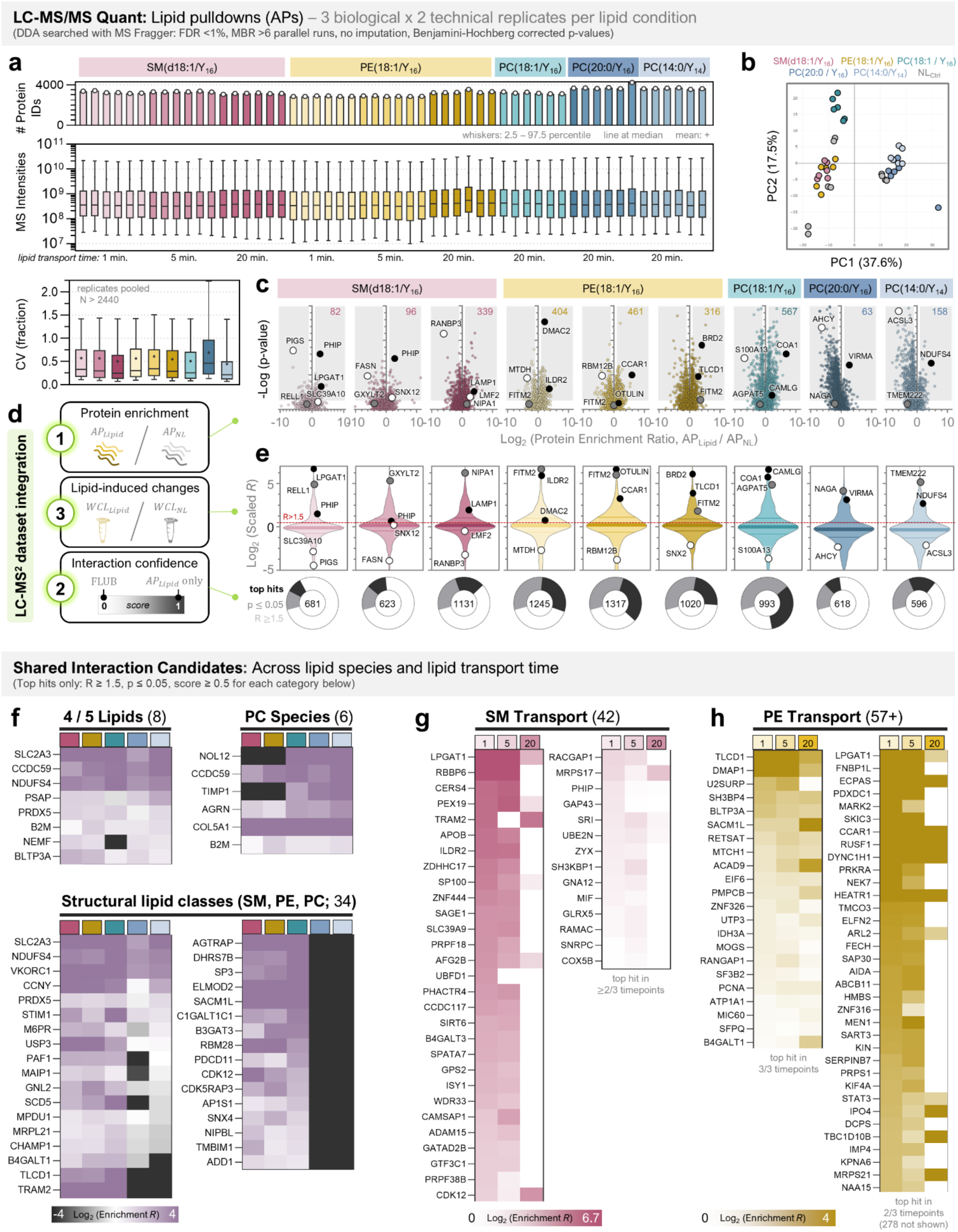
Quantitative proteomics analysis of lipid-protein affinity pulldown (LP-AP) samples. **A)** Summary of LC-MS/MS datasets resulting from Lipid_AP_ samples (N=54 total; 5 lipids, 3 biological x2 technical replicates, 3 timepoints for SM and PE). *Top*, the number of high-confidence protein IDs; *middle*, the distribution of MS intensities for quantified proteins (Benjamini-Hochberg corrected FDR < 1%, minimum 2 unique peptides per protein, see **Methods**); *bottom*, the coefficients of variation (CV) for all proteins per condition following averaging of replicates (N>2440 proteins). Data is shown as box-and-whisker plots with line at median, + at mean, whiskers 2.5-97.5%. **B)** PCA plot of Lipid_AP_ replicates as indication of variability. Color key is at *top*; only shown for 20-minute timepoints. **C)** Volcano plots of the mean protein enrichment ratio (prior to WCL scaling) for every condition (lipid + timepoint), with ratio of AP_Lipid_ to AP_NL_ on the *x*-axis and p-value on the *y-*axis. Each dot is a protein; several are labeled by their confidence assignment as in **E** (black = top hit, grey = interaction candidate, white = not an interactor). The number of significantly enriched proteins (*right* shaded region; R≥1.5, p≤0.05) is indicated at the top of each plot. **D)** Schematic summary of the three datasets integrated for the final list of top interaction candidates. First, the enriched proteins as in **C.** Second, proteins are scaled by their WCL abundance as in **E.** Third, proteins are assigned confidence scores based on NL control analyses as in **Extended Data Figure 3**. **E)** *Top*, Violin plots showing the distribution of individual protein abundances after scaling to WCL (total protein abundance in starting sample). The same proteins are labeled on this plot as in the above volcano plot from **C**, illustrating the relative abundance shifts for different proteins following scaling. The dotted red line indicates the enrichment threshold of R>1.5, which is used for most analyses in this study. *Below*, Donut plots indicating the proportion of enriched proteins (N>618) that are assigned high confidence as top interaction candidates (black; R≥1.5, p≤0.05, score≥0.5). **F)** Heatmaps of top hits shared across lipid species at the 20-minute timepoint. Proteins are divided into 3 categories: shared by 4/5 lipids (N=8), shared by all PC species (N=7), or shared across SM, PE, and PC with (18:1/Y_16_) tails (N=34). Heatmap key is *below*, whereby purple represents enrichment and grey represents depletion from the Lipid_AP_ dataset, plotted on a Log_2_ scale; protein names are at *left.* Note that proteins can be enriched (purple) yet not pass the p-value or scoring filters for the given category. **G-H)** Heatmaps of top hits shared across 1, 5, and 20 minutes (*left-to-right*) of retrograde lipid transport for **G) SM(d18:1/Y_16_)** or **H) PE(18:1/Y_16_)**, plotted as in **F** but with pink (SM) or gold (PE) representing enriched proteins. For SM, only 1 protein is enriched at all 3 timepoints (LPGAT1) and the remaining shown are enriched at 2/3 timepoints (N=41). For PE, 22 proteins are enriched at all 3 timepoints (*left*) and 313 are enriched at 2/3 timepoints (*right*; N=35 are shown, ordered by rank at 1 minute).

**Extended Data Figure 5.**
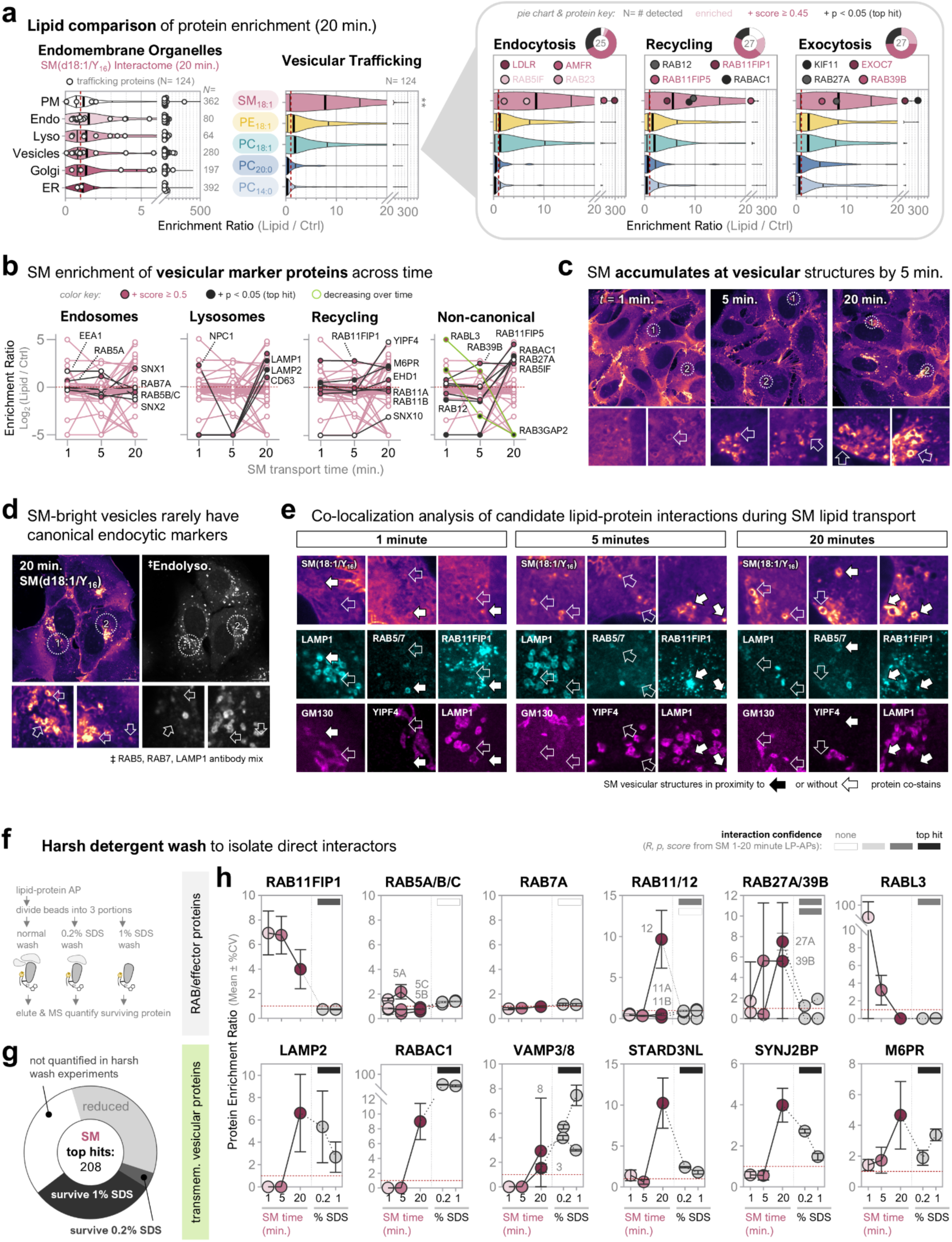
SM exhibits time-specific enrichment of vesicular trafficking components. **A)** Comparisons of endomembrane vesicular trafficking protein enrichment (*x*-axes) in the SM(d18:1/Y_16_) 20-minute interactome compared to the other phospholipids in our study. *Left*, The SM(d18:1/Y_16_) interactome dataset was sorted into endomembrane organelle annotations (*y*-axis) or a function in vesicular trafficking (white dots; N=124); vesicular trafficking proteins are among the most enriched proteins for every endomembrane organelle. *Mid*, The vesicular trafficking protein enrichment (N=124) are compared to the other membrane phospholipid interactomes; SM exhibits higher enrichment of the vesicular trafficking population than all other lipids in our study (**p<0.01 by two-way ANOVA). *Right*, Vesicular trafficking proteins are further categorized by endocytosis (N=37), recycling (N=36), or exocytosis (N=42) and their enrichment is compared across lipid species; for SM, 4 individual proteins are shown per category, colored by interaction confidence (black= top hit; red = R>1.5, score >0.05, p>0.05; light pink = R>1.5). Donut plots above are colored the same, showing the proportion of enriched and top hits out of N = total # of proteins detected in the SM dataset. **B)** Line plots showing the temporal enrichment of vesicular proteins by SM(d18:1/Y_16_), filtered for common “marker” proteins of vesicular compartments (N=45 total; each line is a protein, each dot is the mean protein enrichment value (*y*) at the given timepoint (*x*)). Individual proteins are highlighted in black lines for endosomes, lysosomes, recycling compartments, and non-canonical RAB-associated proteins, colored by interaction confidence (black filled circle = top hit; *key* at top). Lime green lines indicate the only 2 proteins that decrease linearly over time. **C)** Lipid imaging of SM(d18:1/Y_16_) transport in U2OS cells at 1, 5, and 20 minutes post-lipid pulse (150X spinning disc confocal; field of view is 100×100µm). ROIs highlight regions that have SM signal at vesicular structures, indicated by arrows. **D)** Lipid imaging of SM at 20 minutes, comparing localization to co-stains for the canonical endolysosomal pathway (antibodies against RAB5, RAB7, and LAMP1 used in the same channel). SM vesicular structures (arrows), of varying intensity, rarely show co-localization or proximity with RAB5/7/LAMP1. Error bars and ROIs are 10µm. **E)** Imaging validation of SM interactome data, comparing lipid-protein proximity from 1, 5, to 20 minutes of SM retrograde transport. Representative 10×10µm ROIs are shown for each co-labeling condition: LAMP1 with GM130 (Golgi), RAB5/7 (mix) with YIPF4, and RAB11FIP1 with LAMP1. Arrows indicate SM vesicular structures that are distinct from protein co-labels (hollow arrows) or show proximal signal (white arrows). RAB11FIP1 was the only protein to exhibit clear proximity to SM-enriched vesicles at all timepoints. **F)** Schematic of the harsh wash experiment, whereby increasing % SDS is used prior to elution to wash away indirect protein interactions. **G)** Donut plot indicating the proportion of top hits for SM(d18:1/Y_16_) that are maintained following harsh washes; from the 208 top hits identified in the original 20 min. interactome, 144 were detected in the harsh wash experiments (N=2 replicates) and 50% of those (N=72) survived SDS treatment. **H)** Temporal enrichment and harsh wash data for individual vesicular proteins, sorted by RABs/RAB effector proteins (*top* row) and transmembrane vesicular proteins (*bottom* row). Protein enrichment (ratio to control; mean ± %CV) is plotted on the *y*-axis with condition (timepoint or % SDS) on *x.* Interaction confidence from the SM interactomes is indicated as a colored box in the upper-left corner of each graph (*key* at top). Red dotted lines indicate R=1 for visualization.

**Extended Data Figure 6.**
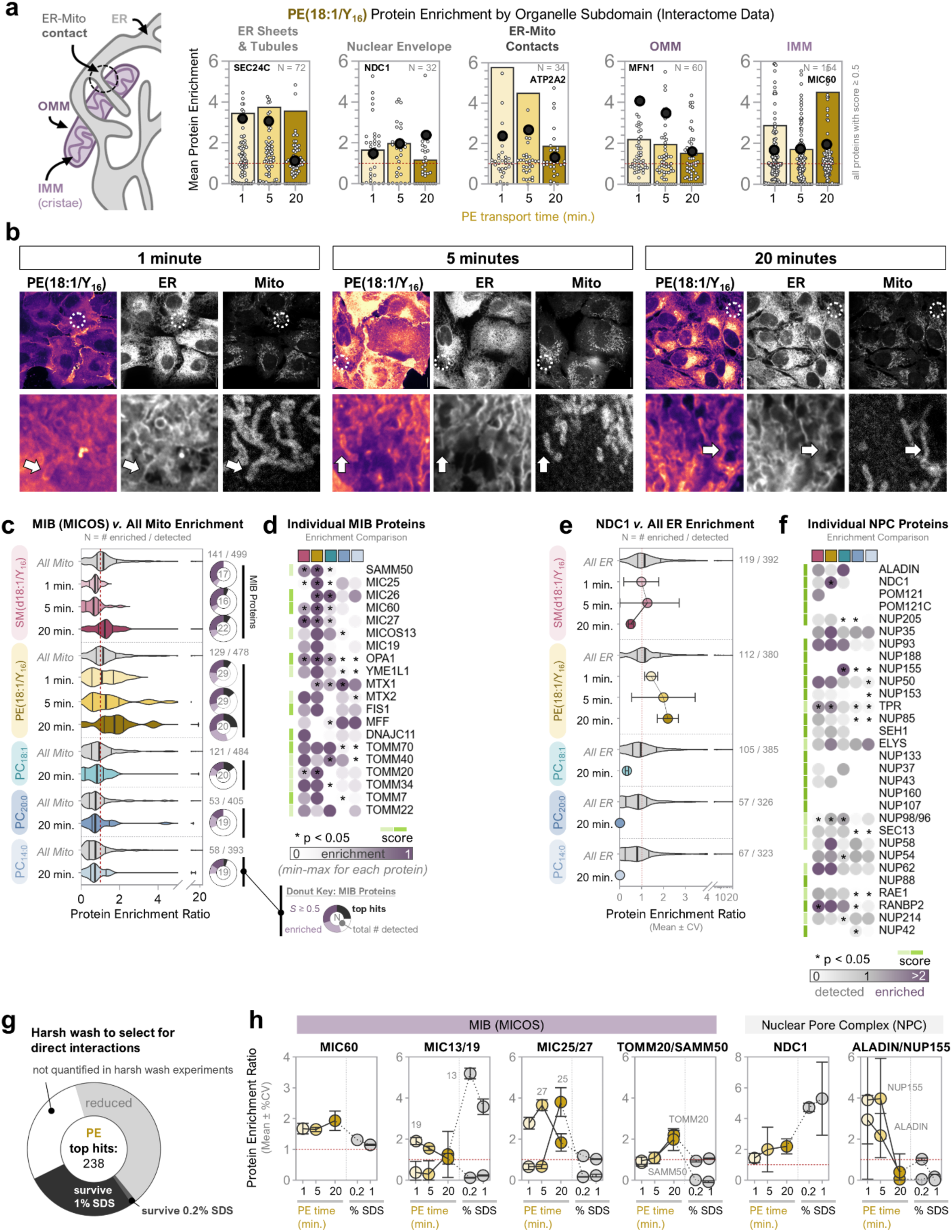
PE engages MIC60 at the mitochondria and NDC1 at the nuclear envelope at the expense of other lipids and proteins. **A)** Comparisons of ER (N=72), nuclear envelope (N=32), ER-mitochondrial contact site (N=34), and mitochondrial outer (N=60) or inner (N=154) membrane protein enrichment (y-axis) by PE(18:1/Y_16_). A representative top-hit interaction candidate from each category is shown as black dots and labeled at the top of each graph; dots are individual proteins, bars at mean. There is an increase in the relative abundance of PE-protein interactions at the inner mitochondrial membrane over time, with concomitant decreases in ER-mitochondrial contact site and OMM proteins, while ER protein interactions are largely stable. **B)** Lipid imaging analysis of PE(18:1/Y_16_) at ER and mitochondria during lipid transport in U2OS cells (150X spinning disc confocal; ROIs are 10×10µm). Arrows indicate PE signal at mitochondrial subdomains, showing a shift from OMM to IMM signal. **C)** Violin plots of mitochondrial inter-membrane bridging space (MIB, contains MICOS; N>16) proteins compared to all other mitochondrial proteins (N>393, grey violins) for each lipid AP in our study (*y-*axis; bold line at median, thin lines at quartiles). A donut plot summarizing the MIB lipid-protein interactions is shown to the *right* or each violin, indicating the proportion of top hits (black) out of the total # of detected MIB proteins for each lipid (*key* at *bottom right*). PE enriches for more MIB proteins than all other lipids tested, and MIB enrichment is elevated above all other mitochondrial proteins. **D)** Heatmap displaying the relative enrichment of MIB complex members across lipids (scaled min-max for each protein, *key* at *bottom*). The confidence score for each protein is indicated in lime green at *left*, * p<0.05, protein names at *right*, and each lipid interactome in a separate column (20 minutes post-lipid loading; color label at *top* colored as in **C**). PE enriches for the majority of MIB proteins. **E)** Violin plots of ER proteins (N>323, grey violins) compared to the mean NDC1 enrichment (error bars ± %CV) for each lipid in our study. PE is the only lipid to exhibit significant enrichment of NDC1, which is among the most enriched ER proteins in the interactome. **F)** Heatmap displaying the relative enrichment of nuclear pore complex (NPC) members across lipids, organized as for **D.** Protein enrichment is shown on a scale from 0 (white; detected yet depleted) to 1 (grey; detected yet not enriched) to >2 (purple; enriched) to highlight the depletion of most NPC proteins. **G)** Donut plot indicating the proportion of top hits for PE(18:1/Y_16_) that are maintained following harsh washes; from the 238 top hits identified in the original 20 min. interactome, 172 were detected in the harsh wash experiments (N=2 replicates) and 40% of those (N=68) survived SDS treatment. **H)** Temporal enrichment and harsh wash data for individual MIB or NPC proteins. Protein enrichment (ratio to control; mean ± %CV) is plotted on the *y*-axis with condition (timepoint or % SDS) on *x.* Red dotted lines indicate R=1 for visualization.

**Extended Data Figure 7.**
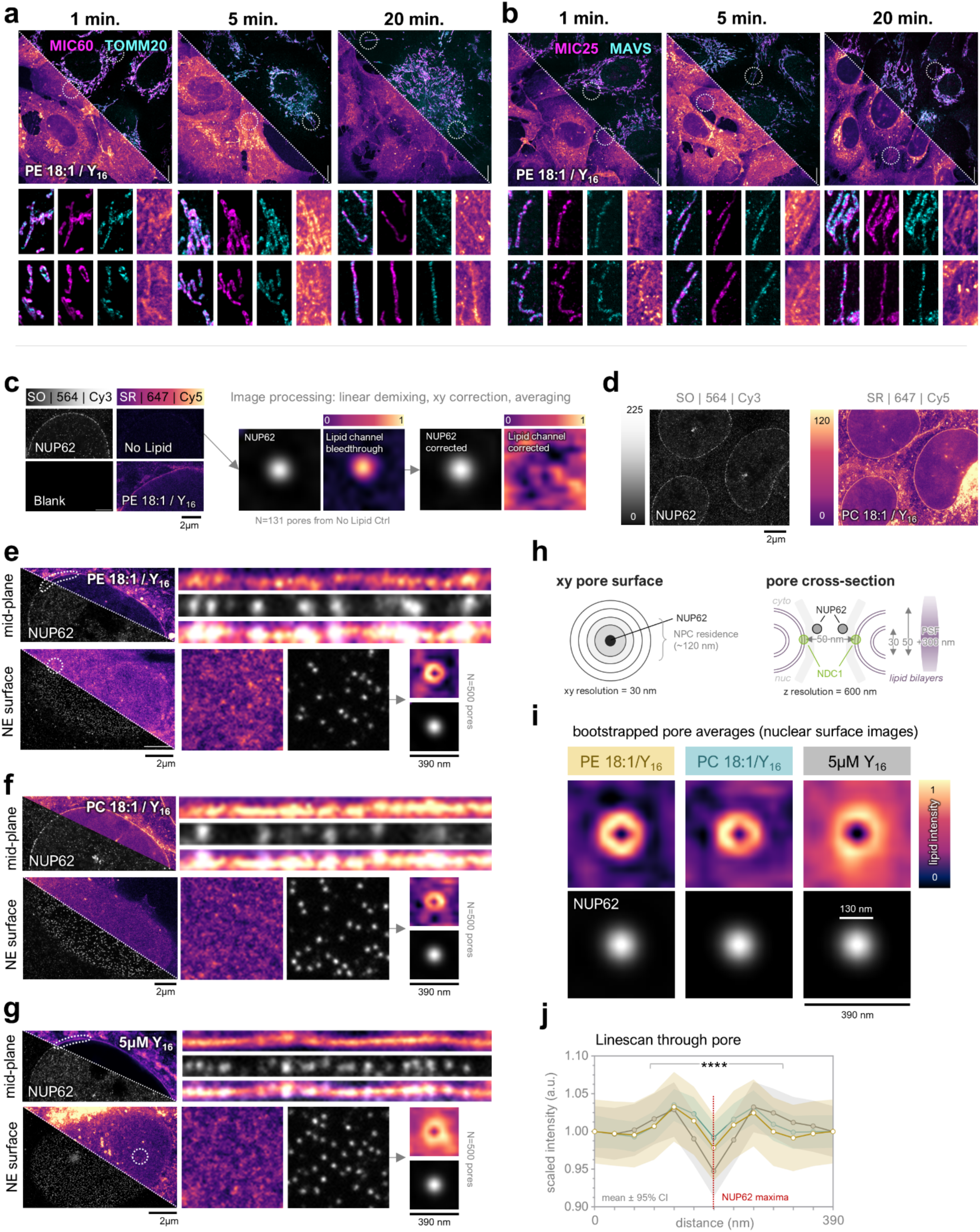
Super-resolution microscopy analyses of PE localization to mitochondrial cristae and nuclear pores. **A)** Lipid imaging of PE(18:1/Y_16_) at 1, 5, and 20 minutes post-lipid loading in U2OS cells co-labeled with antibodies against endogenous MIC60 (magenta) and Tomm20 (cyan), lipid-protein interaction candidates localizing to the MICOS or outer mitochondrial membranes, respectively. Scale bars are 10µm, ROIs are of individual mitochondria resolvable in a single z-plane. *Below*, violin plots of the relative lipid intensity (a.u.) corresponding to areas of each protein co-stain versus the total mitochondrial signal (*p<0.05, **p<0.01, ***p<0.001 by two-tailed student’s t-tests). **B)** Lipid imaging for PE(18:1/Y_16_) versus MIC25 (MICOS) and MAVS (outer mitochondrial membrane), organized as in **A.** **C)** *Left,* Representative 2D STED control images showing minimal bleedthrough between channels. The fluorophore (SO or SR), excitation laser (594 or 647), and emission filter (Cy3 or Cy5) are indicated for each channel at top; all images are shown on the sample intensity scale per channel. *Right*, example of the image processing workflow using the control NUP62 bleedthrough images to compensate for bleedthrough into the lipid channel and align pores using Gaussian means. The “lipid bleedthrough” channel intensity is shown on a min-max scale to visualize bleedthrough, which was <7% of signal from the “normal” lipid images (as visualized at *left*). **D)** Example image from the STED dataset, showing a mid-plane slice of three nuclei labeled for PC(18:1/Y_16_) and NUP62. Channel intensity scales are shown as gradient bars to the left of each image. **E)** Example mid-plane (*top*) and nuclear surface (*bottom*) STED images for PE(18:1/Y_16_), with a linearized ROI of the nuclear rim and a 2×2µm pixel area ROI of the nuclear surface, respectively (1px = 30nm). For the nuclear surface, an example 390×390nm averaged pore sampling is shown (N=500 NUP62 puncta averaged to generate the pore patch). **F-G)** Example STED images for **F)** PC(18:1/Y_16_) and **G)** 5µM Y_16_, organized as for **E. H)** Cartoon schematics of nuclear pore size and structure^96^ influencing STED analysis in the xy (*left*) and xz planes (*right*). The point spread function (PSF) of our STED microscope in the z dimension is ∼300nm, which collapses the lipid signal from the double-membrane structure of the nuclear pore into a single plane. NDC1 is localized at the apex of the outer pore membrane while NUP62 is in the center of the pore. **I)** Averaged lipid signal patches (390×390nm) for all nuclear surface images in our STED datasets (N>2000 pores from N>6 nuclei per lipid condition). Shown are the bootstrap maps for each lipid condition (pure averages are in Figure 4; see **Methods**). **J)** Linescan quantification of lipid signal through the nuclear pore from averaged pore images as in **I** (N>2000 pores from N>6 nuclei). Plotted is the mean lipid intensity at each pixel distance from the NUP62 maxima, with 95% confidence intervals (CI) as the shaded regions (PE=yellow; PC=teal; Y16=grey as in **I**; ****p<0.0001 by ANOVA).

**Extended Data Figure 8.**
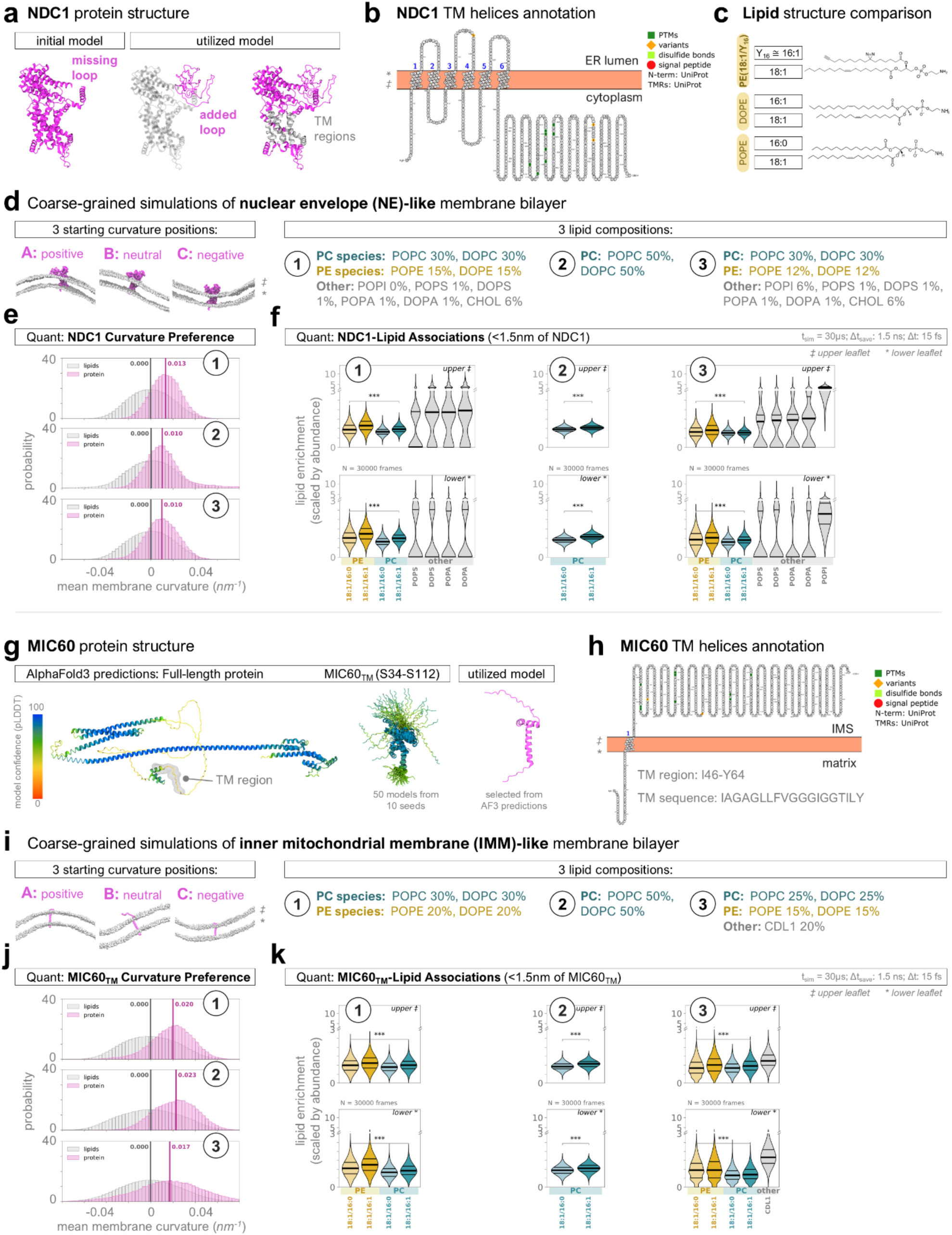
Molecular dynamics (MD) simulations of trans-bilayer lipid interactions with NDC1 and MIC60 in nuclear pore and mitochondrial cristae-like buckled membranes. **A)** NDC1 was modeled from the published structure of the dilated human nuclear pore^92^ (PDB ID: 7R5J, chain JA) with a missing loop (residues G391-P515) was added with Modeller^126^. **B)** Transmembrane (TM) regions and membrane orientation were verified using Protter^127^. Shown is the predicted C- to N-terminus orientation of NDC1 across the nuclear envelope membrane (‡ is the outer/cytoplasmic leaflet, * is the inner/luminal leaflet). **C)** Comparison of PE lipid structures used in simulations (common native species DOPE: PE(18:1/16:1) and POPE: PE(18:1/16:0)) against the bifunctional PE(18:1/Y_16_). The bifunctional Y_16_ behaves similarly to palmitoleic acid (16:1)^14,128^. **D)** Simulated membranes for NDC1 were NE-like in terms of curvature (r=50nm) and lipid compositions (see **Methods** for citations). *Left*, NDC1 was placed at three starting positions relative to the membrane buckle (NDC1 in *magenta*, lipid heads in *grey*; ‡ is the outer/cytoplasmic leaflet, * is the inner/luminal leaflet). *Right,* three different lipid compositions were assessed: 1) NE-like without PI, which we observed to outcompete PE for NDC1 interactions but was not included in our interactome analyses; 2) no PE present; and 3) NE-like including PI. **E)** Quantification of NDC1 (magenta) localization along the buckled membrane, shown as probability histograms relative to lipids (grey). A histogram is shown for each simulated lipid composition (1, 2, 3 as in **D**). Bold lines and associated values indicate the median (N=30,000; bin size = 0.003 *nm^-1^*). NDC1 curvature preference decreases by 3% upon loss of PE. **F)** Lipid association with NDC1 for each simulated lipid composition (1, 2, 3 as in **D** above) at both the upper (*top*, ‡) and lower (*bottom*, *) leaflets of the bilayer, shown as violin plots (bold line is median, thin lines quartiles) of the fraction of lipid species (scaled by molar abundance) <1.5nm from the protein in each frame of the simulation (N=30,000). PE species always outcompete PC species for NDC1 interactions, more pronounced at the lower leaflet (***p<0.001 by one-way ANOVA with Brown-Forsythe multiple comparisons test). **G)** MIC60 structural predictions by AlphaFold3 of the full-length human protein (*left*, color scale from N- to C-terminus) and MIC60_TM_ (S34-S112; *right*, color scale by model confidence). Published human structures were unavailable, but other studies of the MICOS complex^93,106^ and the Protter annotations (**H)** were referenced for reasonability of structural predictions. The utilized MIC60_TM_ model was randomly selected from the 50 AlphaFold3 predictions. **H)** MIC60 transmembrane (TM) helices and membrane orientation as predicted by Protter (‡ is the outer/IMS leaflet, * is the inner/matrix leaflet). I) Simulated membranes for MIC60_TM_ were IMM-like in terms of curvature (r=30nm) and lipid compositions (see **Methods** for citations), whereby PE, PC, and cardiolipin (CDL) are the primary lipid components. *Left*, MIC60_TM_ was placed at three starting positions relative to the membrane buckle (MIC60_TM_ in *magenta*, lipid heads in *grey*; ‡ is the outer/IMS leaflet, * is the inner/matrix leaflet). *Right,* three different lipid compositions were assessed: 1) without CDL, which is known to be the primary driver of membrane curvature at the mitochondria and was not included in our interactome analyses; 2) no PE present; and 3) IMM-like including CDL. **J)** Quantification of MIC60_TM_ (magenta) localization along the buckled membrane, shown as probability histograms relative to lipids (grey). A histogram is shown for each simulated lipid composition (1, 2, 3 as in **D**). Bold lines and associated values indicate the median. MIC60_TM_ exhibits higher curvature preference in the presence of PE without CDL, further shifting by 3% upon loss of PE. **K)** Lipid association with MIC60_TM_ for each simulated lipid composition (1, 2, 3 as in **D** above) at both the upper (*top*, ‡) and lower (*bottom*, *) leaflets of the bilayer, shown as violin plots (bold line is median, thin lines quartiles) of the fraction of lipid species (scaled by molar abundance) <1.5nm from the protein in each frame of the simulation (N=30,000). PE species always outcompete PC species for MIC60_TM_ interactions, more pronounced at the lower leaflet (***p<0.001 by one-way ANOVA with Brown-Forsythe multiple comparisons test).

**Extended Data Figure 9.**
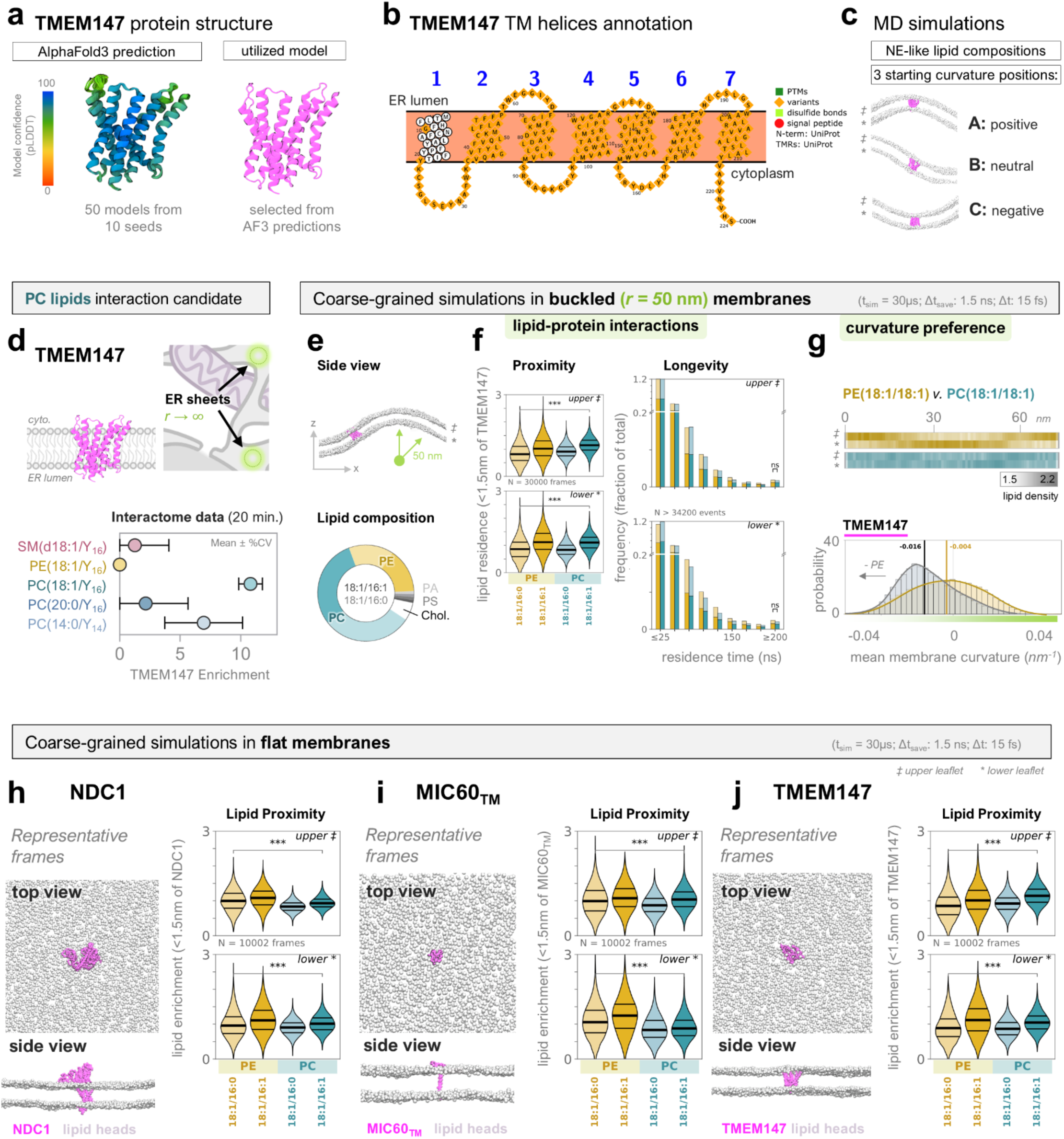
Comparison of lipid interactions with the ER sheet protein TMEM147, NDC1, and MIC60_TM_ in curved versus flattened membrane simulations. **A)** TMEM147 was modeled using AlphaFold3 (color scale is model confidence). **B)** Transmembrane (TM) regions and membrane orientation were verified using Protter^127^. Shown is the predicted C- to N-terminus orientation of TMEM147 across the ER membrane (‡ is the outer/cytoplasmic leaflet, * is the inner/luminal leaflet). **C)** Simulated membranes for TMEM147 followed the same scheme as for NDC1, using ER/NE-like curvature (r=50nm) and lipid compositions (3 total; see **Extended Data Figure 8**). TMEM147 was placed at three starting positions relative to the membrane buckle (NDC1 in *magenta*, lipid heads in *grey*; ‡ is the outer/cytoplasmic leaflet, * is the inner/luminal leaflet). **D)** Structure of TMEM147 used in MD simulations (*left*), cartoon of ER sheet localization indicating the mean radius of curvature at resident membrane domains (*right*), and the mean protein enrichment from the lipid pulldowns (*below,* as in Figure 2 and Figure 5). **E)** *Top,* Side view of the buckled membrane (r=50 nm; representative timeframe of the xz-axis; lipid headgroups = grey, proteins = magenta). *Bottom,* Donut plot of the lipid composition (simulation 3 as in **Extended Data Figure 8**). Each lipid class includes equimolar ratios of palmitoleic (16:0/18:1) and di-oleic (18:1/18:1) derivatives; PC (teal) and PE (yellow) are highlighted in bright colors to match the interactome datasets. Simulations were 30µsec (Δt = 15 fsec). Each replicate started with the proteins initially positioned at different curvatures (positive, neutral, negative; N=3). **F)** Quantification of protein interactions with PE (yellow) and PC (teal) lipid species, shown for both the upper (‡, *top*) and lower (*, *bottom*) leaflets of the bilayer. Lipid proximity is the residence fraction of each lipid species <1.5nm from the protein in each frame (N = 30,000), shown as a violin plot (thick line = median; ***p<0.0001 by Brown-Forsythe multiple comparisons test). Interaction longevity is reported as a histogram of the fraction of lipids remaining <0.6nm from the protein for >1.5nsec (bin size = 25nsec, N>34200 events). PC either outcompetes (upper leaflet) or is very similar to (lower leaflet) PE in proximity to TMEM147, and lipid-protein interactions persist for similar times. **G)** Positioning of lipids (*top,* heatmaps) and proteins (histograms) as a function of membrane curvature. For the heatmaps, the maximum lipid density at each x coordinate (averaged across *y* and time, N=30,000 frames) is shown for both the upper (‡) and lower (*) leaflets of the bilayer, with PE(18:1/18:1) in yellows and PC(18:1/18:1) in teals, plotted on the same scale (*grey* key below). In every simulation, PE enriches at the lower leaflet of positive curvature. For the histograms, the frequency distribution of protein curvature residence (*nm*^-1^, *x*-axis) at every timepoint is shown (N=30,000; bin size = 0.003 *nm^-1^*, median is labeled). TMEM147 exhibits no/neutral curvature preference in the presence of PE (grey). Loss of PE shifts TMEM147 curvature preference to negative, shown as a yellow overlay of the corresponding histogram from simulations containing 0% PE (“-PE”, yellow arrow indicates direction of shift). **H)** Simulations of NDC1 in flat membranes. Representative xy (*top*) and xz (*bottom*) frames are shown, with NDC1 in magenta and lipid headgroups in grey. *Right*, Quantification of NDC1 interactions with PE (yellow) or PC (teal) species at the upper (‡, *top*) and lower (*, *bottom*) leaflets of the bilayer, plotted as in **F**. **I)** Flat membrane simulations for MIC60TM, shown as for NDC1 in **H.** **J)** Flat membrane simulations for TMEM147, shown as for NDC1 in **H.**

## Supplementary information

### Annex: Lipid-Induced Proteome Changes

Using 10% of the starting lysate from every lipid pulldown, we assessed lipid-induced changes to total protein levels by whole-cell proteome analysis (N=4963 proteins, 3 biological x 2 technical replicates; **Supplemental Figure 1, Supplemental Table 6**). These datasets were primarily used for scaling the protein intensity values derived from LC-MS/MS analyses of protein enrichment following lipid-protein affinity pulldown (LP-AP), thus accounting for the technical variability in peptide detection and quantification that results from lipid-protein crosslinking, as described in the main text. In addition, these datasets can provide interesting insights into the bioactive potential of common versus rare membrane lipid species.

In summary, by 20-minutes between 0.8-10.5% of quantified proteins were increased in abundance compared to the no-lipid control condition (2-fold, p<0.05), whereas a greater proportion of 4.1-15.5% were decreased (**Supplemental Figure 1D-E**). The majority of affected proteins changed in only one lipid condition, indicating specific cellular responses to delivery of individual lipids. For all probes, the number of decreased proteins exceeded that of increased proteins, likely reflecting the faster kinetics of protein degradation compared to protein biogenesis^129–131^. Supporting this notion is the observation that many components of the protein degradation machinery change in abundance (e.g., SGTA, UBE4A), including the transmembrane E3 ubiquitin ligase RNF185, which was one of only six proteins with significantly increased levels in all lipid conditions (**Supplemental Figure 1H**). However, we note that many decreased proteins were well-detected in the no lipid controls but poorly detected in the lipid conditions, which inflated fold-change values derived from DDA-MS analysis (**Supplemental Table 6**). Therefore, proteins with decreased levels may also be more prone to loss of detectable peptides following lipid-protein crosslinking in our workflow. Even so, we recently showed that bifunctional lipid crosslinking is only 5% productive in ideal conditions^43^, and any observed technical variability likely only affects a small portion of proteins, as for SM and PE we found a much lower number of affected proteins at earlier timepoints (1 and 5 minutes) (**Supplemental Figure 1I**), in line with the expected timescales of protein biogenesis and degradation.

Delivery of common lipid species affected a higher percentage of the proteome and changes tended to be more pronounced (**Supplemental Figure 1D-F**), suggesting that the biological activity of rare lipid species is limited in the number of possible interactions. Gene ontology (GO) overrepresentation analysis (for SM(d18:1/Y_16_) returned several terms associated with vesicle-mediated transport (e.g. COPII vesicle budding, intra-Golgi transport), reflecting the prominent role of vesicular pathways in intracellular SM transport^10,26,42,58,62^. PE(18:1/Y_16_) affected numerous mitochondrial metabolic processes; in fact, the top decreased protein is the transmembrane protease YME1L1 (**Supplemental Figure 1D**), which functions in mitochondrial homeostasis and is directly regulated by cellular PE levels^64,65^. GO terms returned for PC(18:1/Y_16_) center on translation, ribosome biogenesis, and membrane protein import, in-line with the observation that delivery of this lipid increased the largest number of proteins. General cellular stress responses and lipid biogenesis pathways are not among the overrepresented terms in our datasets; however, many proteins involved in fatty acid remodeling are changed, including increased levels of lipolysis stimulated lipoprotein receptor (LSR), lysophospholipid acyltransferase LPCAT3, phospholipase PLBD2, and sphingolipid desaturase DES1 for SM, PE, PC(18:1/Y_16_) probes, and decreased levels of NDUFAB1, a mitochondrial acyl carrier implicated in lipid homeostasis^132,133^, in all lipid conditions (**Supplemental Figure 1F, H; Supplemental Table 6**). This matches our previous findings that lipid probe delivery does not alter the total native lipidome and that the primary metabolic route for bifunctional lipid probes is acyl chain remodeling^14^. Overall, we find rapid, lipid-specific protein abundance changes that reflect the biological activity of individual lipid species. These data provide the necessary context for detecting bona-fide lipid interactors, experimentally distinguishing lipid-regulated from lipid-interacting proteins.

**Supplemental Figure 1.**
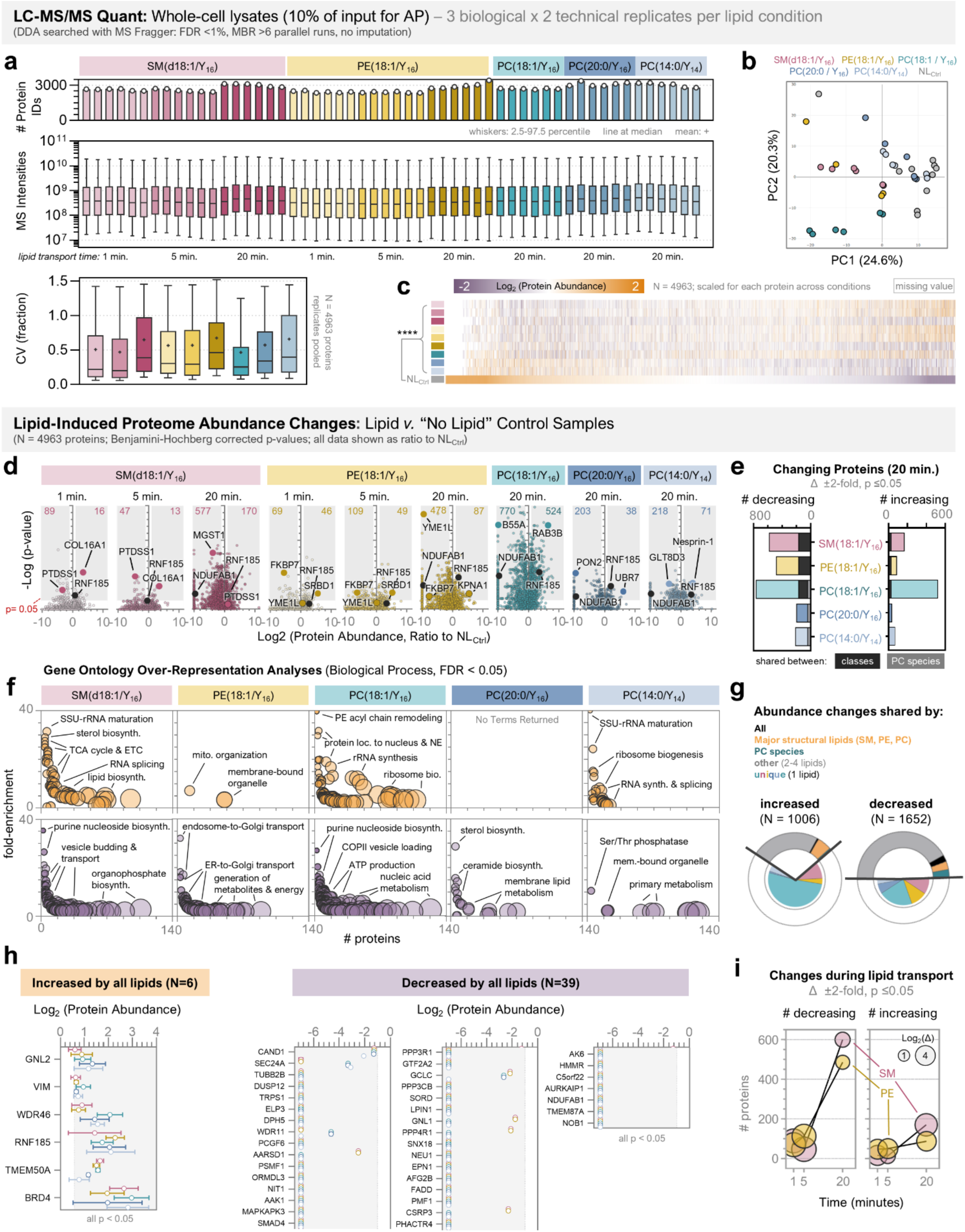
Whole-cell proteome analysis of lipid-induced proteome changes. A) Summary of LC-MS/MS datasets resulting from +lipid whole-cell lysates (Lipid_WCL_) in our study (N=54 total; 5 lipids, 3 biological x2 technical replicates, 3 timepoints for SM and PE). *Top*, the number of high-confidence protein IDs; *middle*, the distribution of MS intensities for quantified proteins (Benjamini-Hochberg corrected FDR < 1%, minimum 2 unique peptides per protein, see **Methods**); *bottom*, the coefficients of variation (CV) for all proteins per condition following averaging of replicates (N=4963 proteins). Data is shown as box-and-whisker plots with line at median, + at mean, whiskers 2.5-97.5%. **B)** PCA plot of Lipid_WCL_ replicates as indication of variability. Color key is at *top*; only shown for 20-minute timepoints. **C)** Heatmap of Lipid_WCL_ protein abundances compared to the NL control (grey, bottom row). Each column is an individual protein (N=4963) scaled min-max across lipid conditions (abundance scale at *top*, lipid species/timepoint indicated by color block at *left* which match labels in **A-B**). Proteins are sorted by relative abundance whereby proteins most abundant in NL controls are at *left* and proteins least abundant in NL controls at *right*. ****p<0.0001 by one- and two-way ANOVA for all comparisons to NL control. **D)** Volcano plots of the mean protein abundance changes for every condition (lipid + timepoint), with ratio of abundance change on the *x*-axis and p-value on the *y-*axis. Each dot is a protein. The number of significantly increasing (*right* shaded region; R≥2, p≤0.05) or decreasing (*left* shaded region; R≤0.5, p≤0.05) proteins is indicated at the top of each plot. Specific proteins are highlighted (black dots = shared by all lipids). **E)** Stacked bar chart depicting the shared (between lipids; *grey*, key *below*) as a portion of the total number of proteins changing in abundance upon delivery of each lipid probe (whole-cell proteomic data filtered for proteins changing in abundance >2-fold, p <0.05, compared to no lipid controls (Δ)). N=3 biological x 2 technical replicates for all datasets in this figure. See also **Supplemental Table 2** for the complete whole-cell proteome data. **F)** Significantly increased (*top,* orange nodes) or decreased (*bottom*, purple nodes) proteins were assessed for over-represented gene ontology terms using PAN-GO^57^ biological process. Data is shown as multi-variable plots whereby the number of proteins assigned to each term is on the *x*-axis, the fold-enrichment compared to the human proteome is on the *y*-axis, and the node size again indicates the number of proteins assigned to the category. A few prevalent examples are labeled; all terms were already filtered by FDR-corrected p-value. **G)** Donut plots of shared abundance changes across lipids, for significantly increased (*left*) or decreased (*right*) proteins at 20 minutes. The outer ring shows the proportions shared by different numbers of lipids, with black = all 5 lipids and white = 1 lipid (unique). For protein changes unique to 1 lipid, the corresponding breakdown across lipid species is shown on the inside as a pie chart, whereby the largest category (e.g., PC(18:1/Y_16_) in teal) indicates the lipid that induces the most unique protein changes. **H)** Individual proteins significantly increased (*left*, orange) or decreased (*right*, purple) by all lipids are shown. For each protein (*y*-axis), the quantified abundance for each lipid (colored as in all other figures) at 20 minutes is shown as mean ± %CV. **I)** Multi-variable plots showing that the number of significantly changing proteins (*y*-axis) increases over time following the initial lipid pulse (*x-*axis), and the number of decreasing proteins (*left*) always exceeds that of increasing proteins (*right*), for SM (*pink*) and PE (*yellow*). The circle diameter indicates the mean effect size.

### Comparing lipid localization by imaging to lipid interactome organelle annotations

Our proteomic assay relies on the selective detection of covalent lipid-protein crosslinks by ratiometric analysis. Crosslinking probabilities increase at sites of elevated local lipid densities e.g., canonical hydrophobic lipid binding pockets (e.g., FABP5 for the rare PCs) or specialized nanoscale lipid environments of transmembrane proteins (e.g., PE-MICOS interactions; **Figure 4**). This contrasts with lipid imaging data, where all lipid-protein contacts contribute to the observed signal, which is an integral of all formed crosslinks. Therefore, we expect only a partial correlation between observed intracellular lipid localization and average organellar protein enrichment among interactor candidates. For example, at 20 minutes SM is primarily localized to the plasma membrane, Golgi, and vesicular compartments (**Figure 2A**). While we find that >50% of the top hits map to these organelles (**Figure 2B**), there is still a good portion assigned to the ER and mitochondria, and these are likely due to selective lipid-protein interactions.

## Materials & Methods

### Cell culture

Adherent monolayer cultures of human osteosarcoma cells (U2OS; originally acquired from DSMZ, ACC785) stably transformed with HaloTag-Sec61β, as previously published^47^, were used for all experiments in this study. Cells were cultured in McCoy’s 5A medium containing GlutaMAX (Gibco, 36600-021) supplemented with 10% fetal bovine serum (FBS; Gibco, 10270-106) and 100 U/ml penicillin–streptomycin (Gibco, 15140-122) at 37 °C and 5% CO_2_. Cells were grown for no more than 10 passages and never allowed to supersede 100% confluency.

For the 1, 5, and 20 minute lipid imaging experiments of PE and SM in Figure 3, cells were additionally stably transformed with SNAP-PDGFR_TM_ into a “Halo-SNAP” double expression line. SNAP-PDGFR_TM_ is a SNAP-tag fast transgene facing the exocytoplasmic side of the plasma membrane via the transmembrane domain of human PDGFR (using Addgene plasmid #182009, gift from Kai Johnsson), enabling use of cell-impermeable SNAPtags that circumvent the endomembrane labeling promiscuity and membrane perturbations caused by intercalated plasma membrane dyes, which are not ideal for lipid imaging. 20 single-cell clones were isolated and ultimately 1 was chosen for experiments based on comparable growth and plasmid expression levels to the parent HaloTag-Sec61β and wildtype U2OS cultures.

### Generation of bifunctional phospholipid probes

All probes used in this study were previously synthesized, characterized, and published by our team in either Iglesias-Artola *et al.* 2025^14^ (18:1/Y_16_ species of SM, PE, and PC) or Lennartz *et al.* 2026^48^ (PC(20:0/Y_16_)), with the exception of the short-chain PC(14:0/Y_14_) which was generated for this study using similar synthetic logic. To enable direct comparisons of minimal changes to individual lipid structures, the probes chosen here always contained the functionalized acyl chain at the *sn-*2 position, with a diazirine at C_7_ and an alkyne on the terminal carbons. The corresponding *sn*-1 acylation (18:1) or headgroup (PC species) was otherwise consistent for direct comparisons of minimal changes to individual lipid structures.

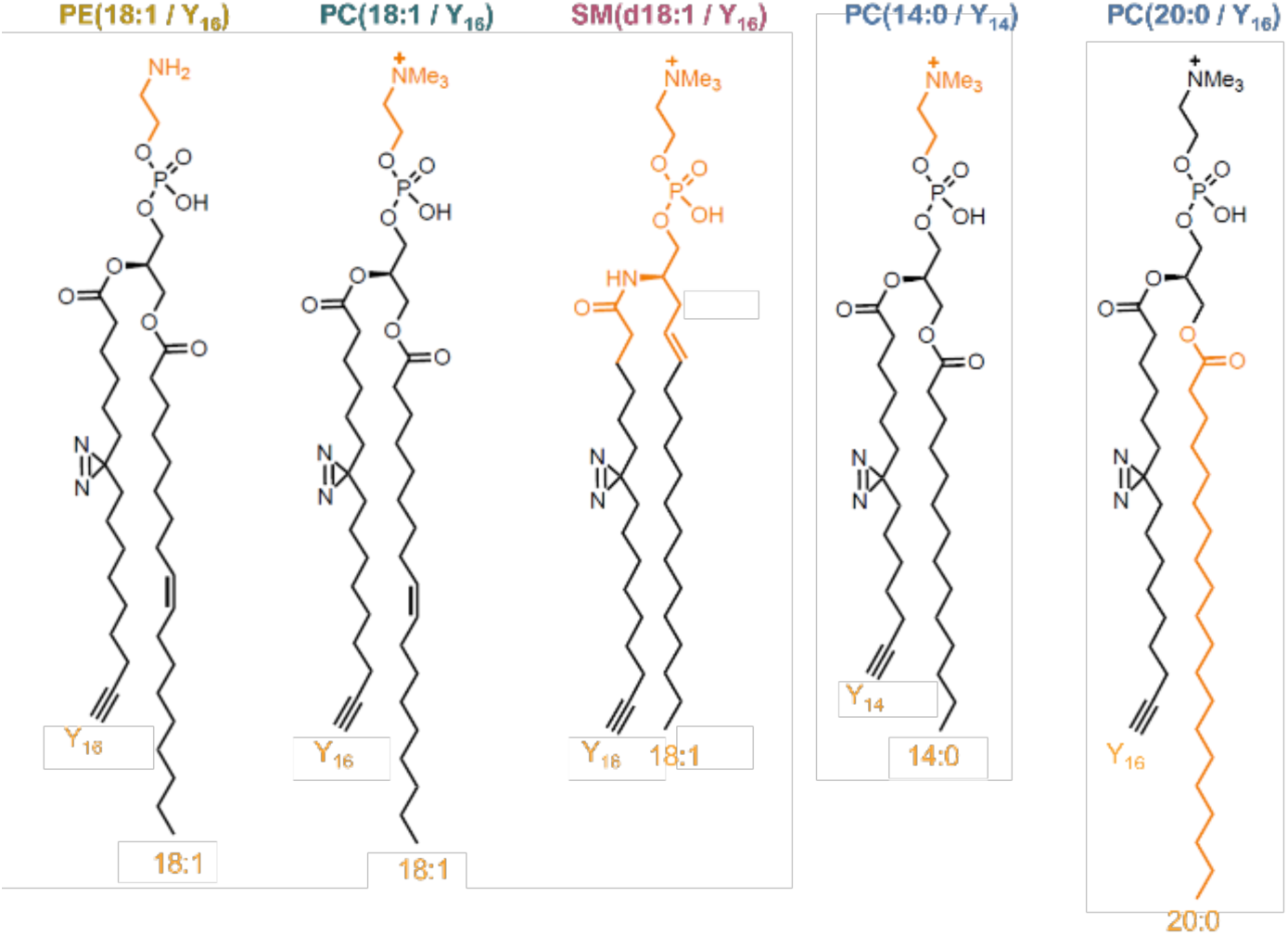

### Lipid pulse-chase using donor liposomes

Bifunctional phospholipids were incorporated into donor liposomes containing 1.5mM bifunctional lipid, 0.75mM POPC (Avanti, 26853-31-6), and 0.75mM cholesterol (Sigma, 57-88-5) in filter-sterilized PBS. Liposomes were filtered to approximately 100nm in diameter by extruding 21x through 0.1-μm polycarbonate membranes (Avanti, 610005-1Ea) using an Avanti Mini-Extruder. Liposomes were stored at 4°C and used within weeks; on the day of use, 0.25mM liposomes were incubated in serum-free media containing 4mM methyl-α-cyclodextrin for >30 minutes at 37°C.

Cells were seeded at a density of 0.8×10^5^ cells/mL and allowed to recover for ∼40 hours or until reaching ∼90% confluency prior to lipid loading. For the loading pulse, cells were washed twice with warm serum-free media and incubated for 1 minute (1-minute timepoints) or 3.5 minutes (5- and 20-minute timepoints) at 37 °C with the liposome/cyclodextrin complex. Time=0 is considered the moment lipid contacts the cells. Cells were then washed twice and returned to free media for the desired chase time (0 for the 1-minute timepoint; 1.5 or 16.5 for the 5- and 20-minute timepoints, respectively). Cells were irradiated using a rapid UV flash (3 seconds; see photoreactor section below) to photoactivate diazirine crosslinking, and either immediately fixed (imaging) or collected in PBS and dropped in liquid N_2_ (LP-AP). A “no lipid” (NL) control was always collected in parallel; all steps were followed except addition of liposomes to the 4mM cyclodextrin/media solution.

### Sample preparation for lipid imaging

All imaging experiments were performed in glass-bottom 96-well plates (Greiner, 655891) so that multiple conditions (e.g., lipid species or timepoints) were monitored in parallel, including one “no UV” control to ensure lipid signal is dependent on photoactivation (**Extended Data Figure 1**). The lipid was always clicked to AlexaFluor 594 picolyl azide (Jena bioscience, CLK-1296-1), which gives the best signal-to-noise in our hands.

Following UV activation, cells were immediately fixed by addition of 4% paraformaldehyde containing 0.5% glutaraldehyde in PBS at 2x volume to each well (working concentration halved to e.g., 2% paraformaldehyde) and left for 20 minutes at room temperature. Cells were permeabilized using 0.1% Triton X-100 in PBS (PBT) for 30 minutes and blocked in 2.5% BSA in PBT for another 30 minutes. Immunostaining was optimized for each target; unless otherwise specified (see antibodies section below), primary antibodies were incubated in blocking buffer overnight at 4°C and Alexa Fluor–conjugated secondary antibodies or HaloTag ligands for 45 minutes at room temperature.

Copper-catalyzed azide–alkyne cycloaddition was always performed after antibody labeling. Fixed cells were washed twice with 100mM HEPES (pH 7.2) and incubated at 37°C with freshly-made click mix: 2µM AlexaFluor 594 picolyl azide, 0.5mM THPTA (Jena bioscience, CLK-1010-1G), 5mM ascorbic acid, and 0.1mM CuSO₄ in 100mM HEPES. Two consecutive click reactions of 22.5 minutes each (45 minutes total click time) were performed, which enhances the labeling efficiency of the self-quenching reaction^47,48^. A final PBT wash for >30 minutes limited background staining; samples were stored in PBS at 4 °C and imaged over subsequent days.

### LED photoreactor design

To build a rapid crosslinking device suitable for large sample sizes, we adapted our previous models^14,47^ to a 15cm diameter array of 324 high-power UV SMD LEDs (λ=365nm; Violumas VS5252C45L9-365). Each LED is equipped with a 90° fused silica lens and positioned 9mm apart, achieving an average of ∼550 mW/cm^2^ illumination with a driving current of 0.35 Amp/LED. The array was set below a metal sample holder to achieve a throw distance of 5mm to the cell surface (including the 1mm thickness of a cell culture plate) that gives relatively uniform illumination across the sample. The entire crosslinking device also contains dimmers for each quadrant, a heat sink (Fischer Elektronik LA-9-200-24) with cooling fans, and a trigger mechanism using a safety cover to prevent dangerous UV exposure. UV rays penetrated both glass and plastic with nearly equal efficiency (measured via luxmeter). The sample stage can be fitted with different inserts to accommodate different sizes of cell culture dishes, and UV activation times can be varied.

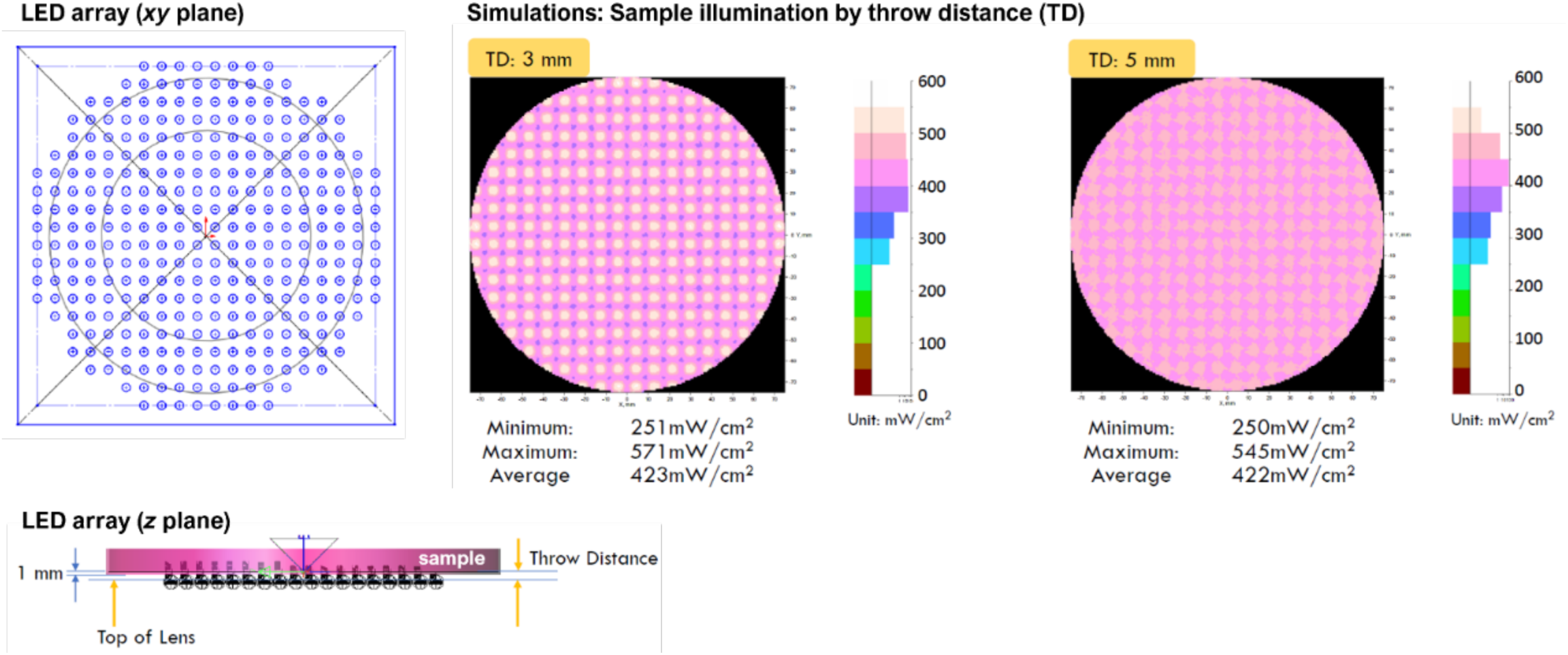

### Spinning-disc confocal microscopy for lipid imaging

Images were acquired using an Olympus IX85 EVIDENT IR Cutter SpinSR spinning disc confocal microscope equipped with an Olympus UPlan N-TIRF 150x (1.45NA) oil objective, Hamamatsu ORCA-Fusion BT Digital CMOS camera, and a Yokogawa CSU-W1 containing both a 50µm and Nikon SoRA spinning disc. The SoRA disc was used to amplify signal but the additional zoom lens for super-resolution capability was not included. All samples were imaged with 0.3µm z-stacks throughout the sample depth, with consistent laser powers and exposure times across experiments, using the Olympus cellSens software (4.3.1) and a channel illumination order of λ = 640, 561, 488, to 405 nm (DPSS lasers). The lipid was always labeled with AF594 and imaged in the 561 channel.

### Lipidomics analysis of bifunctional lipid metabolism

Assessment of SM, PE, and PC 18:1/Y_16_ species conversions was performed using ultra high-resolution mass spectrometry from 4 to 1600 minutes post-lipid delivery in U2OS cells, as reported in Iglesias-Artola *et al.* 2025^14^. For this study, we used these previously published datasets to constrain LP-AP timepoint selection, choosing pulse-chase times whereby no more than ∼10% of the original probe had been metabolized.

### Lipid-protein affinity pulldown (LP-AP)

LP-AP samples were collected from 90-100% confluent 10cm dishes. Before photoactivation, cells were washed 2x with warmed sterile PBS and left in warm PBS for the UV irradiation as described above. Cells were quickly rinsed and scraped into ice-cold PBS, pelleted by centrifugation at 800 x g for 3 minutes at 4°C, and snap-frozen in liquid nitrogen prior to storage at -80°C. Note that only ∼1-3% of the lipidome is exchanged for the bifunctional probe during the loading pulse^14^, which is further distributed throughout the organelle network, and the diazirine forms productive peptide crosslinks at only ∼5% efficiency in ideal conditions, as we recently quantified^43^. Therefore, the following LP-AP protocol and subsequent peptide extraction for MS analysis were optimized to reduce processing steps and time wherever possible to maximize sample yield. The lysis conditions were optimized to retain protein-protein interactions (not crosslinked) in addition to lipid-protein conjugates to boost bioinformatic analysis of proteomic datasets, adapted from previously published interactome studies by the author^134,135^ and group of Dr. Ileana Cristea (Princeton, USA)^50,136,137^.

To account for the variability inherent to membrane biochemistry and the background of a biotin click reaction, a parallel “no lipid” (NL) control was always collected and analyzed in parallel every time the pulldown was performed. We intentionally chose to not use a + lipid / -UV condition as a “control”, as this condition would capture lipid-protein interactions whose stability does not require diazirine crosslinking. In fact, there are several methods where non-covalent protein-lipid complexes are co-isolated for identifying lipid-protein interactions, e.g., for native MS^38^ or for more recently the Lipid-trap technique^37^.

Frozen pellets were resuspended in 300µl of lysis buffer (20 mM HEPES-KOH pH 7.4, 110 mM KOAc, 2 mM MgCl₂, 0.1% Tween-20, 0.6% Triton X-100, 10 µM ZnCl₂, 10 µM CaCl₂, 200 mM NaCl) supplemented with HALT protease/phosphatase inhibitor (Thermo 78438) and nuclease (Pierce, 88700) and incubated on ice for 10 minutes. Samples were then briefly homogenized using a Polytron (8 sec.) and 10% of the lysate was set aside for whole-cell proteome analysis. The remaining 270µl was clicked to biotin picolyl-azide (Jena, CLK-1167) via two consecutive 22.5-minute incubations at 37°C in a 1X working concentration of freshly-made 5X concentrated click mix: 0.74mM biotin picolyl-azide, 1.85mM TCEP, 1.85mM TBTA, and 1.85mM CuSO₄. Use of TBTA/TCEP in our lysate conditions achieved more robust and specific biotin labeling than the THPTA/Ascorbic acid used for lipid imaging (**Extended Data Figure 2**), in agreement with other studies^138^. HEPES as the lysis buffer base was essential (Tris-based buffers confound CuAAC reactions). Insoluble lysate and/or post-click content was retained in the sample; quantifying NL control background is thus critical.

Biotinylated proteins were enriched using Pierce streptavidin-coated magnetic beads (Thermo, 88816). Beads were pre-washed in lysis buffer without enzymes, incubated with clicked lysates for 45 min at room temperature in a sample rotator, washed five times in total with lysis buffer (without enzymes), and eluted by low-pH (Pierce AP elution buffer (pH2)) in a 47 °C shaker for 10 minutes at 800rpm. Eluates were neutralized with Tris base and processed for proteomic analysis; following peptide digest and extraction, the entire protocol yielded enough material for ∼25 LC-MS/MS injections (∼40µg), depending on the lipid.

### Peptide preparation for mass spectrometry (MS) analysis

Whole-cell lysates and LP-AP eluates were prepared for MS analysis using a modified S-Trap (Protifi, C002-MICRO) peptide preparation protocol, optimized for membrane proteins as previously reported in Cook *et al.* 2022^51^. MS-grade reagents and hardware were used throughout. Samples were first brought to 5% SDS by volume, then reduced and alkylated using 25mM TCEP (Thermo, 77720) and 50mM chloroacetamide (Sigma, 22790) for 20 minutes at 70°C. Samples were acidified to 1.2% phosphoric acid prior to binding to the S-Trap columns using 100mM TEAB in 90% LC-MS grade methanol (pH7.5). Columns were washed a total of 5 times at 10,000 x g with the binding buffer. Trypsinization was performed using Promega Platinum (VA9000) at 0.75µg per sample (∼1:30 w/w ratio) in 25mM TEAB for 2.5 hours at 47°C. Peptides were eluted with consecutive 10,000 x g spins in 25mM TEAB, 0.2% aqueous formic acid, and 50% acetonitrile containing 0.2% formic acid for hydrophobic peptides. Peptides were dried in a SpeedVac for 75 minutes at room temperature, then resuspended in 5% formic acid to a final concentration of ∼400 ng/µl for LC-MS/MS analysis.

### LC-MS/MS acquisition and analysis

1.5µg of peptides in 5% formic acid were loaded onto an Aurora Elite C18 UHPLC emitter column with a built-in heater (15cm x 75µm; IonOptics, AUR4-15075C18-XT) using a Thermo Fisher Dionex UltiMate 3000 RSLCnano (UHPLC) system, and separated via a two-stepped linear 90-minute acetonitrile gradient (0-17% until minute 60, 17-35% until minute 80, up to 100% until minute 90). Samples were analyzed by Top20 data-dependent MS analysis (DDA) in a Thermo QExactive Orbitrap mass spectrometer, operating at high MS1 resolution (Rs _m/z 200_

= 120,000; AGC target value of 3e6, max injection time 100ms) within a scan range of 300-2000 *m/z* and using multi-step collision energies (20, 27, 30) for MS2 fragmentation (resolution 60,000, 1e5 AGC, 150ms max injection time, 1.6 *m/z* isolation window). LC-MS/MS methods were managed by Chromeleon and Xcalibur software (Thermo). Blank injections were included to ensure minimal crosstalk between samples and no lipid (NL) controls were always monitored in the same block as the parallel lipid conditions. Polysiloxane (*m/z* 445.12003) was used as lock masses. Two technical replicates were acquired for each sample.

### Spectral searching, filtering, and statistics

All MS raw data used in this manuscript (AP, WCL, harsh wash experiments; N=154 files) were batch-searched altogether using MSFragger 4.0 (Philosopher 5.1.0, IonQuant 1.10.12). Spectra were searched against a concatenated target–decoy database (human, Mycoplasma, and common contaminants) using tryptic specificity (≤2 missed cleavages), peptide mass range of 400-4,000 Da., precursor and fragment mass tolerances of ±15 ppm, cysteine carbamidomethylation as a fixed modification, and methionine oxidation and protein N-terminal acetylation as variable modifications (maximum three variable modifications per peptide). Matched spectra were validated by Percolator-based PSM validation and ProteinProphet protein inference in Philosopher (FDR <1%). Label-free protein quantification (LFQ) was performed with IonQuant; minimal match-between-runs (MBR) was utilized (MBR required ≥6 due to the 6 replicates per condition; *m/z* tolerance 10 ppm, RT tolerance 0.4 min), ultimately applied to <1.35% of peptides across all samples. Protein intensity values (not MaxLFQ) were used for further analysis due to the expected biological variability of missing values in AP versus WCL samples. Further statistical analysis via FragPipe-Analyst^139^ was performed separately for each sample group (AP or WCL), comparing lipid versus NL control conditions with no normalization, minimum imputation, and Benjamini-Hochberg corrected p-values.

Protein intensity values (MSFragger) were exported to Excel for further processing. Each replicate was internally mean-normalized to account for loading/total protein content variability using the mean MS1 intensities before calculating the average protein abundance across replicates for each condition (including standard deviation and coefficient of variation, % CV). Abundance ratios of lipid versus “no lipid” (NL) controls were calculated for each 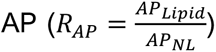 and 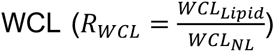. These ratios were paired with the corresponding p-values from FragPipe-Analyst. To account for missing values in the NL controls, simple imputation with a fixed value of either 0.2 for 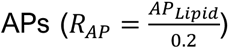 or 0.25 for 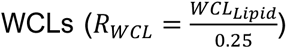 was used to retain lipid-specific proteins in the analysis; these values were roughly the 10^th^ percentile of non-zero protein abundances across the NL samples following mean normalization. Finally, the scaled enrichment ratio 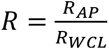 was calculated for each protein in each lipid condition and joined with scores and other annotations in Excel.

### Interaction confidence scoring with false lipid-interacting <u>u</u>nspecific <u>background (FLUB)</u>

Confidence scores (*S*) were generated based on information from no lipid (NL) control datasets (N=14 total samples from 7 biological x2 technical replicates corresponding to 5×20-minute and 2×5-minute conditions; see **Extended Data Figure 3**). Following LC-MS/MS analysis, batch spectral searching by MSFragger (where all lipid and control samples were searched in tandem), and internal mean normalization for each sample, as described above, the average protein intensities from *AP_NL_* and *WCL_NL_* were compared as an enrichment ratio 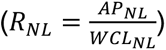 per protein and the associated p-values were calculated (two-tailed and heteroscedastic two-sample t-test). Scores were assigned from 0 (lowest) to 1 (highest confidence) based on the relative enrichment across all proteins, with the following rules:

i) proteins that were significantly enriched by the assay alone (*R_NL_*≥1.5, p≤0.05), without lipid probe, receive *S*=0;
ii) proteins that were never detected in *AP_NL_* samples, but were present in either the *WCL_NL_* or *AP_Lipid_*, represent best-case scenario and receive *S*=1;
iii) all proteins that were significantly depleted by the assay (*R_NL_*≤0.75, p≤0.05) receive scores scaled between 0.5≤*S<*1 by *R_NL_*, whereby the lowest abundance proteins are closest to *S*=1;
iv) all other proteins that did not have significant p-values (p>0.05) and exhibited variable enrichment received scores scaled between 0<*S*<0.5 by *R_NL_*.

This process was performed for each technical replicate pair of *AP_NL_* and *WCL_NL_* samples separately (N=7 scores for each protein) and averaged into the final score reported in main figures and Supplementary Tables 2-3. If a protein had >3 incidences of *S*=0, the averaged score was set to 0.

For defining the final list of proteins considered FLUB (N=716), ratios and p-values were calculated with all NL control replicates combined (**Extended Data Figure 3C-E, Supplemental Table 1**). CRAPome^54^ comparisons were performed by uploading the Uniprot accession codes for all FLUB proteins to the CRAPome repository (https://reprint-apms.org) to assess frequency in the category “H. sapiens -- Proximity Dependent Biotinylation”.

### Annotation of protein features

Gene ontology overrepresentation tests were performed using PAN-GO^57^, comparing against the human proteome with Fisher’s exact FDR-corrected p-values. Any terms reported in this study were filtered for significance (≥2-fold enrichment, p≤0.05). Transmembrane proteins were annotated based on the presence of transmembrane domains in Uniprot^140^. Membrane-associated proteins were annotated based on inclusion in the gene ontology parent category GO0016020 (“membrane” cellular component). For organelle annotations, we manually curated localization assignments by combining information from the following resources: i) the Human Protein Atlas^4,141^, filtered for “approved” reliability at minimum; ii) past peer-reviewed datasets from Jean Beltran *et al.* 2016/2018^142,143^ and Hoyer *et al.* 2024^144^; iii) manual validation using literature search for the top enriched proteins. Proteins were allowed multiple organelle assignments with sufficient evidence, accounting for multi-localized proteins, or were given the descriptor “unknown” if evidence was insufficient. For functional annotations used for categorical comparisons (e.g., vesicular trafficking proteins in Figure 3, OMM and IMM proteins in Figure 4), assignments were given based on keyword searches in Uniprot (e.g., "KW-1000" for OMM and "KW-0999" for IMM; reviewed and human only) and/or manual literature review. For protein complexes, the entire database for human cells was downloaded (https://mips.helmholtz-muenchen.de/corum/) from CORUM^145^.

The resulting “KCC_Proteome-Annotations” are included in descriptor columns in all supplemental tables and provided separately for the human proteome (N=20439) in **Supplemental Table 4** as a resource for future studies.

### Assessing lipid-protein interaction stability by harsh wash

Prior to LP-AP elution, each sample was divided into three equal portions while still on the beads: 1) washed normally, 2) washed with 0.2% SDS, or 3) washed with 1% SDS supplemented in wash buffer for 5 minutes at room temperature. Samples were then eluted, prepared for LC-MS/MS, and analyzed in technical duplicate as for all other LP-AP samples. Following the raw data search using MSFragger (also included in the batch search with all other samples), each replicate was scaled internally to the mean to normalize for loading/total protein content differences between samples. The resulting protein values can be compared against each other directly for each lipid set (Normal, 0.2% SDS, or 1% SDS), given that the three samples were derived from the same starting material. See all replicate data in **Supplemental Table 5**.

### Analysis of lipid signal with mitochondrial proteins

Lipid intensities and Pearson’s correlations with co-labeled mitochondrial proteins were calculated using FIJI Macros in ImageJ. For lipid intensities, multi-point masks were generated from thresholded binary images of each protein co-stain (e.g., MIC60 in 488 and TOMM20 in 405), selecting for the brightest puncta (∼1% of total signal). The mean lipid intensity corresponding to each co-stain’s mask was then calculated for every image in the dataset. For Pearson’s correlations, linearized images of individual mitochondria were manually generated using the segmented line tool (width=15px), picking 10-15 individually resolvable mitochondria in a single z-plane per image. Pearson’s R values for each lipid-protein pair were calculated for each mitochondria using the Coloc2 plugin.

### Stimulated emission depletion (STED) super-resolution microscopy

Two-dimensional STED images were acquired using a commercial confocal infinity-line STED microscope (Abberior Instruments) equipped with a pulsed laser excitation (640nm, 560nm, 460nm, 40MHz), beam scanning module (line frequency 3kHz), single photon APD detectors, and a 100X oil objective (Olympus, 1.49 NA). Lipids were always clicked with a StarRed picolyl-azide (Abberior; excitation λ=640nm / emission range λ=685-70nm), and NUP62 was labeled with a StarOrange secondary antibody (Abberior; excitation λ=561nm / emission range λ=605-50nm). For both channels, depletion was achieved with 50% power of a 775 nm, 40MHz pulsed laser (Katana HP, 1W, 1ns pulse duration, NKT Photonics). Images were acquired at a pixel size of 30nm with a pinhole size of 45µm (xy PSF = ∼60nm), using the Imspector software (v.16.2.8415).

### STED image analysis of lipid signal at nuclear pores

STED experiments included individual channel controls to account for bleedthrough, xy alignment controls using Tetraspecks (100nm; Thermo, T7279), and “no lipid” conditions to assess non-specific fluorophore click background. Images were processed in ImageJ and Python. For nuclear rim images, linearized ROIs were generated using the segmented line tool (width=8px) for display in figures; no additional image processing occurred. For nuclear envelope surface images, we performed a particle averaging analysis to resolve lipid signal around the nuclear pore using custom python scripts integrated in Napari written by ChatGPT 5.5. The NUP62 signal was used to detect NUP particles. Particles were filtered based on area and minimum separation distance (area=5-15px; min separation distance=250nm) and particles at nuclear edge and beyond were excluded using user-defined masks. Particles were aligned using 2D Gaussian fits of the NUP62 signal, generating a 390 x 390 nm stack of aligned pores containing the NUP96 and the bifunctional lipid channels. The lipid and protein signal of all aligned particles (N ∼3000 pores per lipid probe) was then averaged. To correct the bleed-through of the NUP62 signal into the lipid channel, we used control images of NUP62 alone (StarOrange), from which we determined the XY shift between the two channels and the bleed-through by minimizing the signal in the lipid channel. Once the correction parameters were determined they were fixed and used for correction of all lipid samples.

For data interpretation including cartoon representations of NDC1 and NUP62 orientations in nuclear pore architecture, we relied on several excellent reviews, especially Petrovic *et al.* 2026^96^ and Rothballer and Kutay 2013^114^.

### Molecular Dynamics (MD) simulations

All simulations were performed with the Martini 2.2 coarse-grained force field^146^, which has been employed to recreate the curvature preferences of lipids^147,148^ and proteins^149^. The workflow comprised protein model preparation, membrane construction, a series of equilibration steps, membrane buckling, protein insertion, and production runs. Identical simulation parameters were used for buckled and flat-membrane control systems unless noted otherwise. All MD simulations were performed using GROMACS 2024.5^150^ at the CHEM cluster operated at the Max Planck Computing and Data Facility in Garching.

#### Protein structure preparation

Three transmembrane proteins were studied: NDC1, MIC60, and TMEM147. NDC1 is a component of the NPC. The NDC1 structure was taken from the NPC dilated state^92^ (PDB ID: 7R5J, chain JA), with a missing loop between residues G391 and P515 added with Modeller^126^ (**Extended Data Figure 8A**). MIC60 is part of the MICOS complex; the full-length AlphaFold model did not predict the transmembrane segment as a helix (UniProt Q16891). A truncated model from residues S34 to S112 (MIC60_TM_) was generated, including part of the protein facing the matrix and the intermembrane space. Fifty models were created from 10 random seeds using AlphaFold3^151^. The transmembrane segment (residues I46 to Y64) was predicted as a helix in all truncated models, and one model was selected randomly for simulations (**Extended Data Figure 8G**). TMEM147 is an ER protein; the sequence was taken from UniProt Q9BVK8 and its structure predicted with AlphaFold3. Fifty models were made from 10 random seeds, and one model was selected randomly. All predictions had high confidence and were similar (**Extended Data Figure 9A**). Each of these proteins were first protonated at pH 7.0. The protonated structure was then solvated in explicit TIP3P water^152^ and 150 mM NaCl, and energy-minimized using the all-atom CHARMM36m force field^153^ (steepest-descent, 10 kJ mol⁻¹ nm⁻¹). After minimization the structure was converted to a Martini representation with martinize.py^146^ (elastic network with default protein settings) and secondary structure restraints assigned via DSSP^154^.

#### Membrane composition and system building

Buckled and flat membranes were simulated. NDC1 and TMEM147 used endoplasmic-reticulum (ER)–like compositions. MIC60_TM_ used inner-mitochondrial-membrane (IMM)-like compositions. Initially, flat membranes were built with insane.py^155^. Three lipid mixtures were used for the ER– like membranes^1,156,157^ of NDC1 and TMEM147 with initial box size 70nm × 30nm × 18nm: (1) POPC 30%, DOPC 30%, POPE 15%, DOPE 15%, POPS 1%, DOPS 1%, POPA 1%, DOPA 1%, CHOL 6%, (2) POPC 50%, DOPC 50%, and (3) POPC 30%, DOPC 30%, POPE 12%, DOPE 12%, POPI 6%, POPS 1%, DOPS 1%, POPA 1%, DOPA 1%, CHOL 6%. For MIC60_TM_, IMM-like compositions^1,108,158^ for an initial box size 50nm × 20nm × 18nm were: (1) POPC 30%, DOPC 30%, POPE 20%, DOPE 20%, (2) POPC 50%, DOPC 50%, and (3) POPC 25%, DOPC 25%, POPE 15%, DOPE 15%, CDL1 20%. All systems were solvated with Martini water containing 10% anti-freeze WF particles and 150 mM NaCl. Buckled membrane curvature was based on prior structural work of the nuclear pore^92,96^ and mitochondrial cristae^93,94,98,108^, choosing r=50nm and r=30nm, respectively.

#### Simulation parameters and equilibration

Non-bonded interactions used the Verlet cutoff scheme. The neighbor list radius was 1.35 nm, updated every 20 steps. Electrostatic interactions used the Reaction-Field method^159^ (*ε_r_* = 15, *ε_rf_* = ∞). Potentials were truncated at 1.1 nm. An energy minimization was performed with the steepest descent algorithm. The maximum force tolerance was 100 kJ mol^-1^ nm^-1^. The systems were equilibrated for 2 ns in the NVT ensemble at 310 K with the velocity-rescaling thermostat^160^ (*τ_t_* = 0.2 ps). Then, equilibration for 5 ns in the NPT ensemble at 310 K was performed. Target pressure was set to 1.0 bar the with Berendsen barostat^161^ (*τ_p_* = 12 ps). The x/y dimensions were fixed and the z-axis relaxed (*β_zz_* = 3.0 × 10^−4^ bar^-1^).

#### Membrane buckling protocol

A compressed state was induced for buckling. A 60 ns simulation at 310 K using the Berendsen barostat^161^ (*τ_p_*: was performed. Lateral load was applied along the x-axis (*P_xx_* = 2.0 bar). The z-axis pressure was 1.0 bar. The y-dimension was fixed. The buckled membrane was relaxed to establish a stable curvature profile in a 60 ns run at 310 K. The x/y box dimensions were fixed. The z-axis relaxed (*P_zz_* = 1.0 bar).

#### Production simulations

Each protein was inserted into the buckled membranes. Three locations were used: positive (crest), neutral (flank), and negative (trough). Overlapping solvent molecules were removed. The system relaxed with a short equilibration following the same protocol as before. A 30 μs production simulation was performed for each configuration. The time step was Δt=15 fs. Temperature was 310 K. The velocity-rescaling thermostat^160^ was used. Pressure was targeted to 1.0 bar along the z-axis with the anisotropic Parrinello-Rahman barostat^162^ (*τ_p_* = 12 ps). The lateral dimensions (x, y) were fixed to keep the membrane buckle.

#### Flat membrane simulations

Flat membranes were simulated as controls. This decoupled membrane curvature from lipid composition. These systems skipped the compression and buckling steps. They used the same force field and composition. The same minimization and equilibration sequence was followed. A 30 μs production simulation (Δt=15 fs) was performed. Temperature was 310 K. Pressure was 1.0 bar. Lateral and perpendicular dimensions were coupled semi-isotropically with the Parrinello-Rahman barostat. This ensured a tensionless, flat bilayer.

### Molecular dynamics simulation analysis

Visual Molecular Dynamics (VMD)^163^ was used for trajectory visualization. All analyses on the coarse-grained trajectories were performed with MDAnalysis^164^ and plotted with Python and Prism. Lipids were tracked through their headgroup phosphate bead (PO4, or PO41 for cardiolipin) and the protein through its backbone beads, with all distances evaluated under periodic boundary conditions. Each observable was computed independently for every replicate over the equilibrated production window (last half of the production runs), and the distributions reported were obtained by pooling or averaging across the replicates of each composition.

#### Leaflet-resolved lipid enrichment

Lipids whose headgroup lay within 1.5 nm of the protein in a given frame were assigned to the protein annular shell. Leaflets were assigned each frame by a graph-based connectivity search over the phosphate beads (MDAnalysis LeafletFinder), taking the two largest connected groups and labelling them upper and lower by their centre-of-mass height. For each leaflet and frame, the shell composition of species *i* was expressed as its mole fraction within the shell, 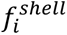, and the bulk composition as its instantaneous mole fraction across the whole leaflet, 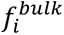. Enrichment was defined as 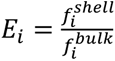, with *E* > 1 indicating recruitment to the shell, *E_i_* < 1 depletion and *E_i_* ≈ 1 a shell composition matching bulk. Per-frame enrichment values were pooled across replicates and shown as violin plots for each species and leaflet.

#### Membrane curvature preference

A continuous membrane surface ℎ(*x*, *y*) was reconstructed for each frame by fitting a two-dimensional truncated Fourier series to the phosphate-bead heights, using the in-plane box dimensions as periodicities^149^. The mean curvature 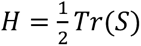 was obtained analytically from the shape operator *S* of the fitted surface and evaluated at the lateral position of the protein centre of mass, yielding a time series of the local curvature sampled by the protein. Curvature values were pooled across replicates and shown as probability-density histograms.

#### Lipid headgroup density

Because the buckle drifted over the trajectory, each frame was first registered to a common reference by fitting a single-mode cosine to the lipid tail-bead heights along the buckling axis and translating the frame so that the ridge was centered laterally and the bilayer midsurface at half the box height. For each species, headgroup positions were then binned onto a fixed grid in the *x* − *y* plane (the plane containing the buckling axis and the bilayer normal) and accumulated over the final 15 µs to give a time-averaged number density projected along *x*. Each map was normalised by the mean density of the membrane phase, defined as the grid region exceeding 5% of the maximum density, giving a dimensionless field equal to 1 at the membrane-average concentration. Normalised maps were averaged across replicates and rendered as heatmaps.

#### Lipid residence times

Residence times were measured per lipid from the minimum distance between its headgroup and the protein, using a dual-cutoff criterion to prevent boundary recrossings from fragmenting a single binding event: a lipid was counted as bound on first entering an inner cutoff (0.6 nm) and as released only on crossing a larger outer cutoff (1.5 nm), with the residence time taken as the elapsed time between these events. Leaflet classification was performed in the same way as for the leaflet-resolved lipid enrichment. Events were pooled across replicates and shown as bar plots for each species and leaflet.

### Protein assessment by SDS-PAGE

During LP-AP optimization, protein samples were analysed by SDS-PAGE followed by Coomassie, Western blotting, or silver stain. Samples were heated at 95°C in LDS buffer (at 1X) for 10 minutes to denature and then loaded on NuPAGE 4-12% Bis-Tris gels (1.5mm; Thermo, NP0322PK2) and separated by weight at 130V for 1.5 hours. Coomassie Brilliant Blue (BioRad, 1610436) was used for ∼30 minutes to stain total protein and washed 4x with water prior to imaging. For Western blotting, gels were transferred to methanol-activated PVDF membranes overnight (∼16 hours) at 30V on-ice (at 4°C), blocked in 2.5% BSA in PBS, stained with primary antibodies for 2 hours in block with 0.2% Tween-20, washed in 0.1% Tween-20 in PBS, and stained with secondary antibodies for 45 minutes in block with 0.2% Tween-20 and 0.01% SDS. Silver staining (0.1% AgNO_3_, 0.02% formaldehyde) was done for 20 minutes after 1 hour of gel fixation (40% ethanol, 10% acetic acid) and 1 minute of sensitization (0.02% Na_2_S_2_O_3_), then washed 5x in water and developed (3% Na_2_CO_3_, 0.05% formaldehyde). Gels and blots were imaged using an Invitrogen iBright.

### Protein co-labeling with antibodies and transgenes

Antibody labeling conditions were optimized for each target protein. HaloTag ligand (diluted to 200nM; JF646, Promega GA1120) was added alongside secondary antibodies to label the ER. SNAP-tag surface ligand (AF488, NEB #S9136) was diluted to 1nM in complete medium and added to live cells for 20 minutes prior to the lipid loading pulse to label the PM. NUP62, MIC60, and MIC25 antibodies showed low efficacy with glutaraldehyde fixation, likely due to the high protein density of their respective locations, so for the respective co-labeling experiments (**Figures 4 and 5**) paraformaldehyde alone was used for fixation.

For lipid imaging experiments: MIC60 at 1:750 (Proteintech, 10179-1-AP), MIC25 at 1:500 (Proteintech, 20639-1-AP), MAVS at 1:750 (Santa Cruz, sc-166583), Tomm20 at 1:500 (Santa Cruz, sc-17764), NUP62 at 1:750 (BD Biosciences, 610497), RAB11FIP1 at 1:250 (Cell Signaling, 12849), LAMP1 (Ms) at 1:750 (Santa Cruz, sc-20011), LAMP1 (Rb) at 1:750 (Cell Signaling, 9091), LAMP2 at 1:750 (Santa Cruz, sc-18822), Rab5 (Ms) at 1:300 (BD Transduction, 610725), Rab5 (Rb) at 1:750 (Cell Signaling, 3547S), Rab7 (Ms) at 1:1000 (Cell Signaling, 95746S), Rab7 (Rb) at 1:750 (Cell Signaling, 9367S), YIPF4 at 1:750 (Proteintech, 15473-1), EEA1 at 1:250 (BD Transduction 610457), SNX1 at 1:750 (Atlas Antibodies, HPA047373), GM130 at 1:750 (BD Transduction, 610822). In Figure 3 and Extended Data Figures 1 & 6, the endolysosome organelle population stain was Rab5, Rab7, and Lamp1 rabbit antibodies mixed at 1:750 each; the mitochondrial organelle population stain was MAVS and Tomm20 at 1:000 each. All AlexaFluor secondary antibodies (H+L Cross-Adsorbed goat-anti-mouse or goat-anti-rabbit fluorophore conjugates, e.g., Thermo A-31556) were used at 1:2000. The Aberrior StarOrange secondary antibody (STORANGE-1050) was used at 1:200.

For Western blotting: GAPDH at 1:2500 (Abcam, ab8245), LaminB1 at 1:500 (Abcam, ab16408), Tomm20 at 1:1000 (Cell Signaling, 42406S), Streptavidin-647 conjugate at 1:2000 (Thermo, S32357).

### Data presentation

All plots were generated using GraphPad Prism 11. Lipid probe chemical structures were generated in ChemDraw. Figure assembly, labeling, and cartoon generation (manual by KCC) was done in Microsoft Powerpoint.

## Notes

### Competing Interest Statement

The authors have declared no competing interest.

### Summary of Updates

A formula in Supplementary Table 2 for top hit filtering contained an error, this has been corrected.

